# Distinct transcriptional profiles of maternal and fetal placental macrophages at term are associated with gravidity

**DOI:** 10.1101/2023.09.25.559419

**Authors:** Nida Ozarslan, Joshua F. Robinson, Sirirak Buarpung, M. Yvonne Kim, Megan R. Ansbro, Jason Akram, Dennis J. Montoya, Moses R. Kamya, Abel Kakuru, Grant Dorsey, Philip J. Rosenthal, Genhong Cheng, Margaret E. Feeney, Susan J. Fisher, Stephanie L. Gaw

**Affiliations:** Division of Maternal-Fetal Medicine, Department of Obstetrics, Gynecology & Reproductive Sciences, University of California, San Francisco (UCSF); San Francisco, CA, USA; Center for Reproductive Sciences and Department of Obstetrics, Gynecology, and Reproductive Sciences, University of California, San Francisco (UCSF); San Francisco, CA, USA; Obstetrics & Gynecology Institute, Cleveland Clinic Foundation; Cleveland, OH, USA; Department of Molecular, Cellular & Developmental Biology, David Geffen School of Medicine, UCLA; Los Angeles, USA; Department of Biochemistry and Molecular Medicine, University of California Davis Health; 2700 Stockton Boulevard, Sacramento, CA, USA; Infectious Diseases Research Collaboration; Uganda; Department of Medicine, Makerere University; Kampala, Uganda; Division of HIV, Global Medicine, and Infectious Diseases, Department of Medicine, University of California, San Francisco; San Francisco, CA, USA; Department of Molecular Immunology and Genetics, University of California, Los Angeles; Los Angeles, CA, USA.; Division of Experimental Medicine, Department of Medicine and Pediatrics, University of California, San Francisco; San Francisco, CA, USA

**Keywords:** pregnancy, placenta, maternal-fetal immunology, reproductive immunology, macrophage, monocyte, Hofbauer cell, gravidity, RNA-seq, transcriptomics, gene expression, innate immunity

## Abstract

Maternal intervillous monocytes (MIMs) and fetal Hofbauer cells (HBCs) are myeloid-derived immune cells at the maternal-fetal interface. Little is known regarding the molecular phenotypes and roles of these distinct monocyte/macrophage populations. Here, we used RNA sequencing to investigate the transcriptional profiles of MIMs and HBCs in six normal term pregnancies. Our analyses revealed distinct transcriptomes of MIMs and HBCs. Genes involved in differentiation and cell organization pathways were more highly expressed in MIMs vs. HBCs. In contrast, HBCs had higher expression of genes involved in inflammatory responses and cell surface receptor signaling. Maternal gravidity influenced monocyte programming, as expression of pro-inflammatory molecules was significantly higher in MIMs from multigravidas compared to primigravidas. In HBCs, multigravidas displayed enrichment of gene pathways involved in cell-cell signaling and differentiation. In summary, our results demonstrated that MIMs and HBCs have highly divergent transcriptional signatures, reflecting their distinct origins, locations, functions, and roles in inflammatory responses. Our data further suggested that maternal gravidity influences the gene signatures of MIMs and HBCs, potentially modulating the interplay between tolerance and trained immunity. The phenomenon of reproductive immune memory may play a novel role in the differential susceptibility of primigravidas to pregnancy complications.

## INTRODUCTION

The placenta is the site of maternal-fetal interactions that mediate nutrient, gas and waste exchange between mother and fetus. It also maintains immune tolerance critical for healthy pregnancy. A chimeric organ comprised of tree-like chorionic villous tissue of embryonic origin, the placenta derives its blood supply through maternal spiral arteries and maternal-fetal exchange occurs in the intervillous spaces and chorionic villi (*1, 2*).

From mid-to late gestation, the most abundant leukocyte in the placenta is the monocyte/macrophage (*3, 4*). Placental macrophages are critical for maintaining the tolerogenic environment that is required for a healthy pregnancy and defending against pathogens (*5–7*). Macrophages at the maternal-fetal interface include three distinct cell types differing in location and origin: (1) maternal decidual macrophages, (2) fetal Hofbauer cells (HBCs), and (3) maternal intervillous monocytes (MIMs) (*8*). Maternal decidual macrophages, located at the placental-uterine interface, are the most well characterized and have been shown to participate in angiogenesis, spiral artery remodeling, trophoblast invasion, apoptotic cell phagocytosis, and immunomodulation during normal pregnancies (*9–11*). Their overactivation can result in pathological conditions such as preeclampsia, fetal growth restriction, and stillbirth (*12*).

Despite their identification over a century ago (*13–15*), little is known regarding the phenotype and function of fetal Hofbauer cells (HBCs). They emerge from the yolk sac as early as the 18^th^ day of gestation(*16, 17*). Later in pregnancy, they differentiate from fetal liver monocytes. HBCs remain at high levels throughout the pregnancy (*18*) as tissue-resident macrophages within the chorionic villous cores (*19*). Data suggest that HBCs participate in the immune response to exogenous pathogens (*20–22*), inflammatory responses to maternal disease (e.g. obesity, preeclampsia, and diabetes (*23–26*)), regulation of nutrient transport, placental vasculogenesis, and angiogenesis (*27, 28*).

In contrast to fetal-derived HBCs, MIMs are maternal bone marrow-derived monocytes in the peripheral circulation that enter and exit the intervillous space through maternal spiral arteries and veins, respectively. MIMs have been shown to exhibit increased influx to the intervillous space during spontaneous labor (*29*). They also play a significant role in systemic and placental inflammatory reactions (*30–32*).

MIMs and HBCs express the classic mononuclear phagocyte cell surface receptors, CD68 and CD14(*33*). There is no consensus on specific markers to distinguish MIMs and HBCs, hence they have been discriminated from one another (and from decidual macrophages) chiefly by their anatomic localization at the maternal-fetal interface (*34–36*). Further investigation into the phenotype and role of these distinct placental mononuclear cells has been hindered by the difficulty of isolating pure populations of MIMs and HBCs, as they are located in close proximity.

Proper control of immune activation states at the maternal-fetal interface is required for successful pregnancies (*37*). M1 macrophages form the first-line in host defense against a variety of bacteria, protozoa, and viruses, as well as in anti-tumor immunity (*38*). In contrast, the M2 subset is characterized by diverse immunosuppressive activity, and includes wound-healing macrophages, IL-10-secreting regulatory macrophages, and tumor-associated macrophages (*39*). To date, there have been limited studies investigating the activation states of placental monocyte/macrophages. Although studies are sparse, HBCs have been shown to play an anti-inflammatory role in the placenta and may contribute to maternal-fetal tolerance. These studies have found that HBCs function predominately in the M2 activation state during normal pregnancy (*18, 28, 40–44*). Dysregulation of M1/M2 phenotypes in the placenta has been associated with pregnancy complications, such as atopic disease, gestational diabetes, preeclampsia, preterm birth, and fetal growth restriction as well as susceptibility to bacterial, protozoal and viral (CMV, HIV, Zika) infections (*6, 15, 26, 45–48*). For example, differential activation of M1/M2 ratios in HBCs has been found to be associated with birth weight in placental malaria (*49*).

Limited investigations have characterized monocyte/macrophage cells of the human placenta. In a targeted investigation, HBCs underwent dynamic changes in RNA expression of M1/M2 markers across gestation (*50*). Recent studies utilizing transcriptomic profiling have used a non-biased approach to characterize monocyte/macrophage populations using targeted (cell-selection) or non-targeted (bulk tissue) approaches (*51*). Targeted isolation of lung alveolar macrophages in different disease states revealed great diversity in macrophage activation states (*52, 53*). Other studies leveraging single cell RNA sequencing (scRNAseq) methods on bulk tissue samples have also shown distinct populations of monocyte/macrophage cell types at the maternal-fetal interface, including cells in the placental villi, chorionic membranes, and the basal plate (*54–57*). These studies demonstrate the breadth of macrophage diversity across gestational ages and clinical states and suggest that mononuclear phagocytes may play a role in pregnancy-related health and complications.

We sought to compare the transcriptional profiles of MIMs and HBCs in normal term pregnancies to better understand the phenotypes of these cells at the maternal-fetal interface and to gain insight into their potential functions. We separated MIMs and HBCs and performed bulk RNAseq to characterize the two populations. We also investigated the influence of maternal gravidity on the activation states of maternal vs. fetal immune cells in term placentas. Together, these data provide a valuable analysis of these two cell populations in normal pregnancy against which alterations that occur in pregnancy complications can be interrogated.

## RESULTS

### Location of MIMs and HBCs at the maternal-fetal interface

We isolated paired MIMs and HBCs from human term placentas. Circulating MIMs are located in the intervillous space (IVS) of the placenta, where maternal blood circulates (arrowhead; **Figure 1**). HBCs (arrow; **Figure 1**) reside in the chorionic floating villi (FV), which consists of an outer syncytiotrophoblast (STB) layer, an inner layer of cytotrophoblasts (CTBs), stroma, and fetal blood vessels. MIMs and HBCs were purified from the villous tissue as described.

**Figure 1.**
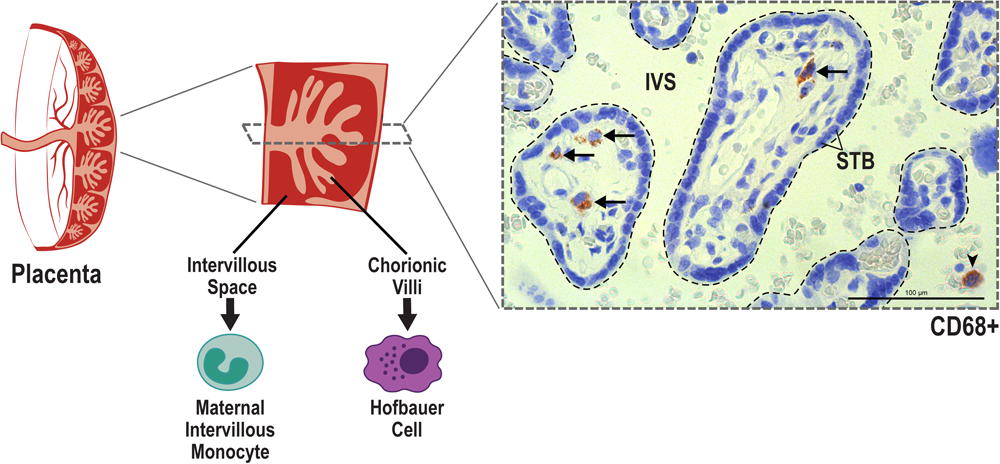
Anatomical localization of maternal intervillous monocytes (MIMs) and fetal Hofbauer cells (HBCs) within the placenta. MIMs and HBCs reside in unique placental compartments within close proximity. HBCs (arrow) originate from the fetal yolk sac and reside in the chorionic villi (encircled with dashed lines). MIMs (arrowhead) are derived from the maternal bone marrow, and ultimately, enter the placenta via the maternal spiral arteries that provide oxygenated blood to the intervillous space (IVS). MIMs and HBCs highly express CD68+ (brown) in histological crosssections of the chorionic villi and intervillous space.

### Differences in gene expression between MIMs and HBCs

We conducted mRNA profiling of MIMs and HBCs. The distribution of average gene counts was similar in the two cell types for all placental samples (**Figure 2A**). The top 2% of abundant genes in MIMs and HBCs (319 genes; dashed line; **Figure 2A**) were significantly enriched for macrophage-progenitor signatures (myeloid CD33+ and monocyte CD14+) as determined by cell type enrichment analysis (**Figure 2B**). MIMs were over-represented in the placenta, suggesting the placental reference database includes a significant contribution from maternal blood. The gene profiles of MIMs vs. HBCs were similar to those of their cognate cells in whole blood. Additionally, these profiles were akin to other organs with high levels of tissue-resident macrophages.

**Figure 2.**
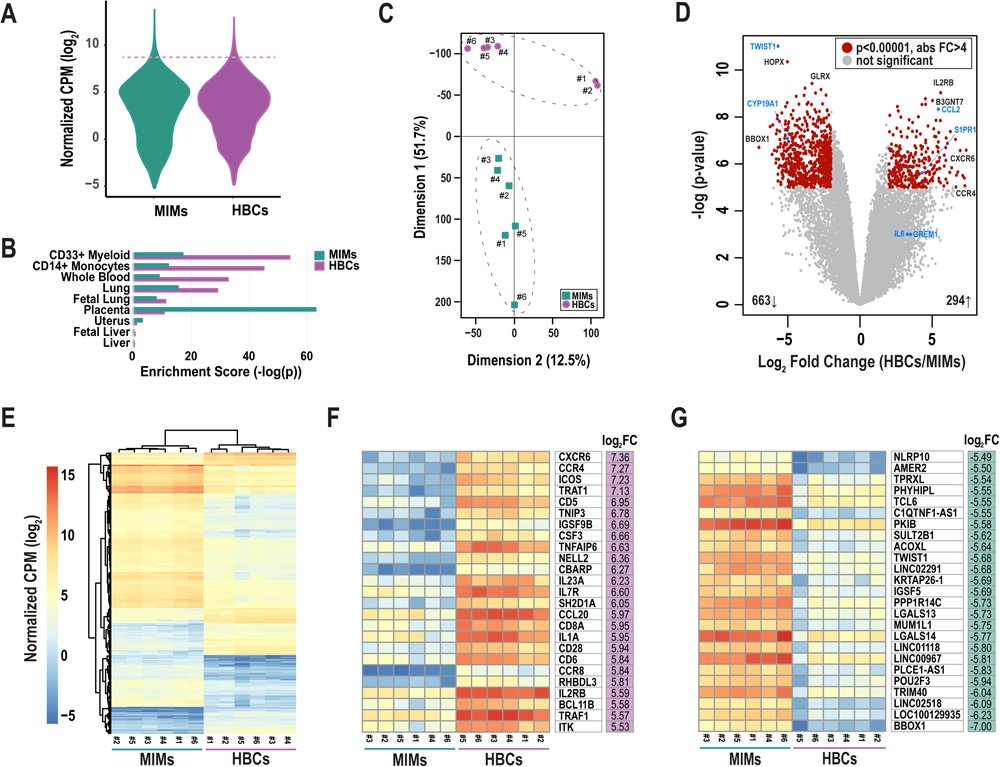
Differentially expressed genes between MIMs and HBCs. (**A**) Distribution of average gene counts (log2 normalized counts per million (CPM)) in MIMs and HBCs. (**B**) Cell/tissue enrichment of the most abundant genes in each cell type (top 2%, dashed line in panel **A**). (**C**) Multidimensional scaling plot of MIM (teal squares) and HBC (purple circles) transcriptomes. Paired MIMs and HBCs denoted by sample IDs. (**D**) Volcano plot displaying significance (negative log p-value) and fold difference in expression (log2). Red dots signify differentially expressed genes between the two cell populations (p<0.00001; absolute FC > 4). Genes labeled in blue were validated via qRT-PCR. (**E**) Hierarchical clustering plot displaying absolute mRNA expression of differentially expressed genes. Expression of the top 25 most up-(**F**) or down-(**G**) regulated genes in terms of fold change (FC) in HBCs vs. MIMs.

Principal component analysis of all transcripts clearly separated MIMs from HBCs with Dimension 1 accounting for 51.7% of the variability (**Figure 2C**). LIMMA identified 976 differentially expressed (**Figure 2D**; red circles) transcripts between the two cell types (p ≤ 0.00001, absolute FC ≥ 4). Within this subset, 663 genes (69%) were upregulated in MIMs whereas 294 genes (31%) were upregulated in HBCs. Hierarchical clustering of differentially expressed genes also highlighted the distinct profiles of these two cell types and confirmed the similarity of samples within each cell type (**Figure 2E**). The 25 most upregulated genes in HBCs vs. MIMs included molecules involved in inflammatory responses (*CXCR6*, *CCR4*, *CCL20*, *CD6*, *CD28*, *IL1A*, *TNFAIP6*; **Figure 2F**). The 25 most upregulated genes in MIMs vs. HBCs included molecules linked to IL-6-regulated cytokine pathways (*NLRP10*, *TWIST1*) and macromolecule metabolism (*PKIB*, *PPP1R14C*, *POU2F3*, *TRIM40*, *TWIST1*; **Figure 2G**). Overall, our analyses demonstrate distinct transcriptomic signatures between these two cell types.

### Validation of genes differentially expressed in MIMs vs. HBCs

To validate the RNA-seq results (Figure 2D, blue marked genes, **Supplemental Table 1**), we used quantitative RT-PCR to assess the expression of selected genes that were highly differentially expressed (*TWIST1, CYP19A1, CCL2, S1PR1;* p<0.00001) and two genes with a more modest difference (*GREM1,* p=0.003; *IL6,* p=0.002). Using a larger sample set, which included additional placentas, we confirmed the expression of *CCL2* and *S1PR1* to be significantly higher in HBCs vs. MIMs and expression of *CYP19A1* and *TWIST1* to be significantly higher in MIMs vs. HBCs (Figure 3A). Expression of *IL6* and *GREM1* was increased in HBCs vs. MIMs. Overall, the magnitude and directionality of expression differences assayed by the two platforms were strongly positively correlated (R^2^=0.8541, Figure 3B).

**Figure 3.**
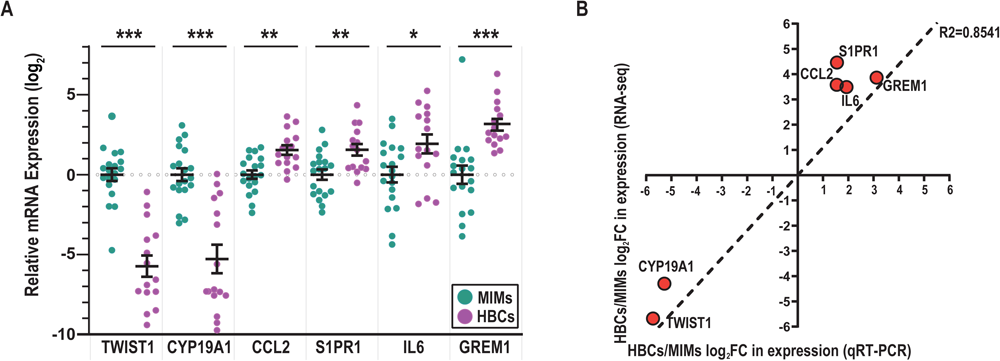
mRNA validation of differentially expressed genes between MIMs and HBCs. (**A**) Relative expression of genes identified to be differentially expressed between HBCs (purple circle) and MIM (teal circle) via qRT-PCR. Expression values (ΔΔCT) were normalized to housekeeping genes (GAPDH, ACTB) and adjusted by the average MIM expression. Asterisks indicate significant differences between MIMs and HBCs (***p<0.0005; **p<0.005; *p<0.05). Bars reflect mean and standard error (SE). (**B**) Correlation plot between RNA-seq and qRT-PCR expression levels of HBC/MIM fold change values in log2 format. Regression line signifies relationship between fold change differences of MIMs and HBCs quantified via RNA-seq vs. qRT-PCR (slope = 1.095; R2=0.8541).

### Mapping differentially expressed genes to Biologic Processes

We evaluated differentially expressed genes between MIMs and HBCs for enrichment of biological processes (Figure 4A). Fifty GO terms were identified as overrepresented (**Supplemental Table 2**) and included processes related to cell motility, adhesion, inflammation, cell death, signal transduction, communication, metabolism, and organization. MIMs had greater expression of genes related to cytoskeleton organization, intracellular signal transduction and cell differentiation. In contrast, HBCs tended toward increased expression of specific biological processes related to inflammation, single organismal cell-cell adhesion and cell surface receptor signaling pathways.

**Figure 4.**
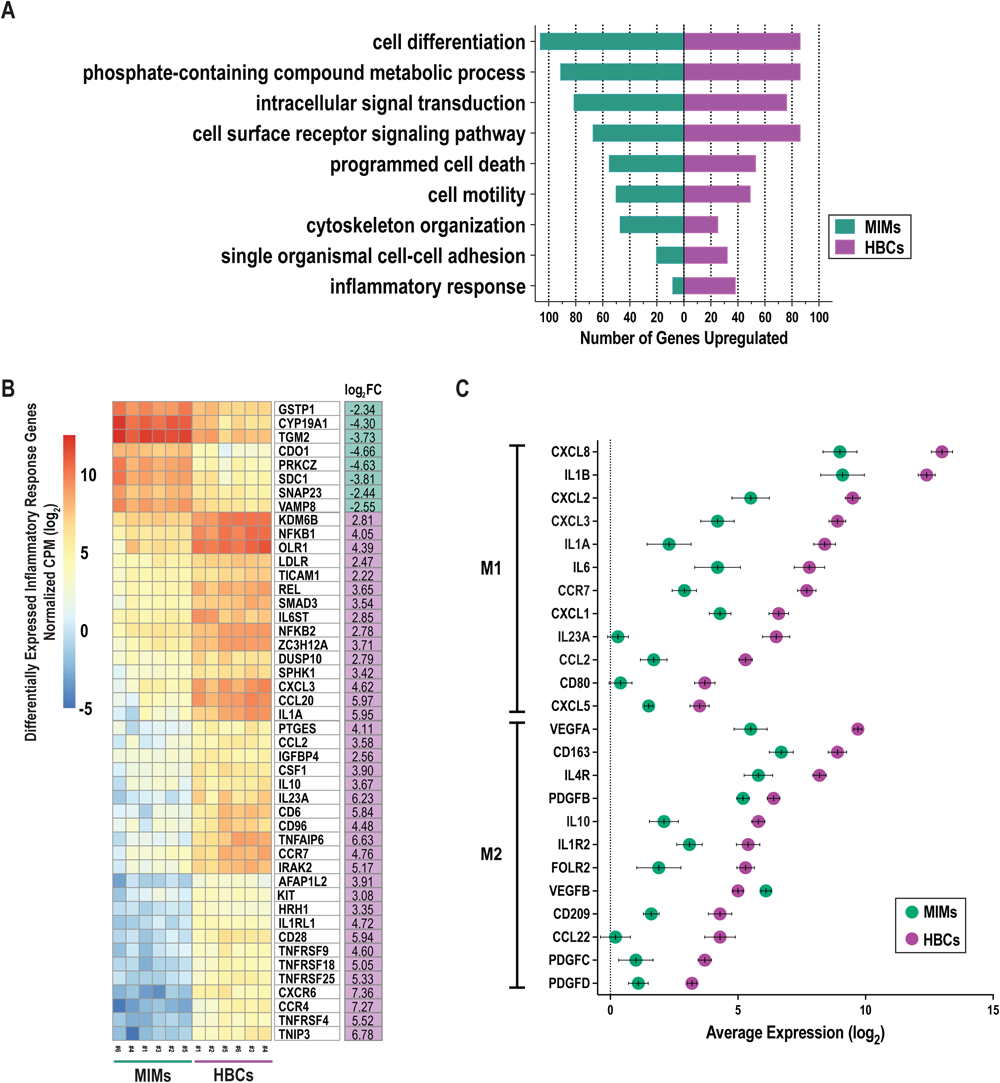
Functional enrichment analysis of differentially expressed genes between MIMs and HBCs. (**A**) Selected enriched biological processes (criteria: p ≤ 0.01, number of differentially expressed genes associated with enriched term ≥ 10) and the number of differentially expressed genes with higher expression in MIMs or HBCs in each category. (**B**) Hierarchical clustering of differentially expressed genes associated with inflammatory response pathway (GO:0006954). (**C**) Relative expression of M1/M2 markers in HBCs vs. MIMs. Panel of 40 genes associated with pro-(M1) and anti-(M2) inflammatory activation states were identified based on current literature. Out of this panel, 24 genes were differentially expressed between MIMs and HBCs (average log2count > 0, p < 0.01 and absolute log2FC > 1). Genes were ordered by decreasing expression in HBCs within each activation state.

Further exploration of a subset of genes involved in the inflammatory response (Figure 4B) confirmed robust differences in gene expression between MIMs and HBCs. Those with known functions in macrophage biology that were more highly expressed in MIMs compared to HBCs included *CYP19A1*, *TGM2*, *SNAP23* and *VAMP8.* Macrophage-associated genes that were more highly expressed in HBCs vs. MIMs were *KDM6B, NFKB1, CXCL3, CCL20, IL1A, CCL2, CSF1, IL10, IL23A, CCR7* and *IRAK2*. Overall, these findings demonstrate that MIMs and HBCs have different transcriptional profiles and suggest potential drivers of differences in the function and behavior of these cells.

### Expression of M1/M2 markers in MIMs and HBCs

We compiled a panel of forty molecules associated with pro-and anti-inflammatory states (M1 vs. M2, respectively) based on published data (*7, 9, 37, 40, 43, 58–71*) (**Supplemental Table 3**). We relaxed our cut-off criteria (p<0.01, absolute log2FC > 1, average log counts > 0) to explore the expression profiles of these markers in MIMs and HBCs. Twenty-six genes were differentially expressed between MIMs and HBCs. Of these, 25 were more highly expressed in HBCs and included M1 and M2 markers (Figure 4C). The average fold-increase of these molecules was 10.5 (range 2.3 to 73.5). Examples of the largest differences included *IL23A*, *IL1A*, *CCL2*, *CCR7*, *CXCL3* and *IL10* (red). *VEGFB* was the only marker that was more highly expressed in MIMs vs. HBCs.

### Gene expression differences by gravidity

We next assessed differences in gene expression in MIMs and HBCs between primigravid and multigravid pregnancies. Gravidity influenced the expression of 120 and 95 genes in MIMs and HBCs, respectively (p<0.05; absolute FC > 2; Figure 5A). Overlap in these subsets was limited to five genes: *C15ORF48, ZNF135, OR2B11, MSC,* and *LINC01291*. Hierarchical clustering showed that in MIMs, 85% (158/186) of the differentially expressed genes were more highly expressed in multigravidas (Figure 5B). In HBCs, differentially expressed genes were equally distributed in the up-and down-regulated categories (50.6% and 49.4%, respectively).

**Figure 5.**
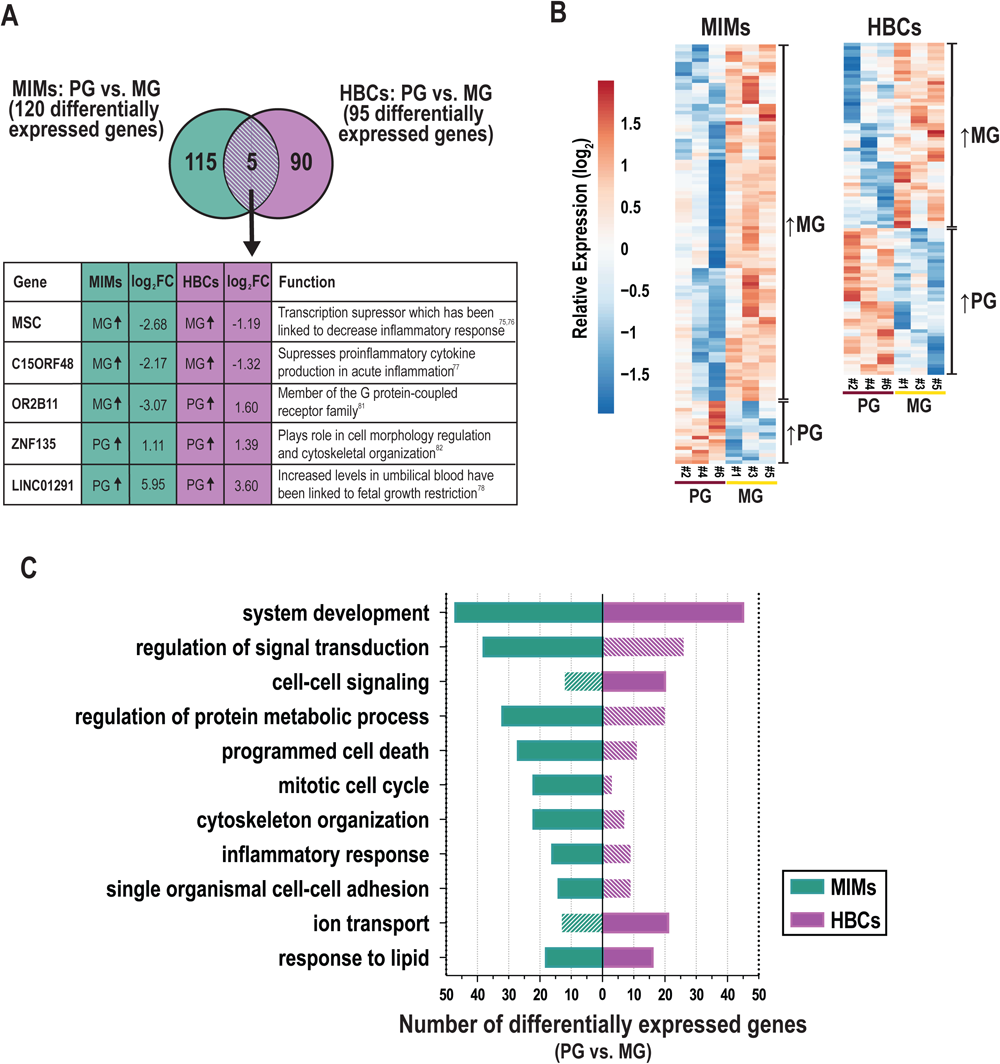
The influence of gravidity on gene expression in MIMs and HBCs. (**A**) Genes identified to be differentially expressed due to gravidity (primigravid or multigravid placentas) in the two cell types (p<0.05; absolute FC > 2). Five differentially expressed genes in common between MIMs and HBCs due to gravidity are shown in the box with their previously described functions(*83–85, 96–98*). (**B**) Relative expression (based on average expression of primigravida) of genes significantly influenced by gravidity in MIMs or HBCs. Hierarchical clustering denote genes higher in multigravida vs. primigravida or vice versa in each cell type. (**C**) Select enriched biological processes associated with gravidity in MIMs or HBCs (p<0.01; number of differentially expressed genes ≥6 in any cell type). Filled bars indicate significance (p<0.01). Hatched bars indicate non-significance.

GO analyses of these data is shown in **Supplemental Table 4**. Figure 5C depicts a portion of the results for each cell type with statistically significant biological processes denoted by solid color bars. System development and response to lipids were the only processes that were significantly differentially expressed in association with gravidity in both cell types. MIM genes that were differentially expressed as a function of gravidity were involved in inflammation (*e.g.,* inflammatory response, response to cytokines); migration/adhesion (*e.g.,* single organismal cell-cell adhesion, leukocyte migration); cell cycle processes (*e.g.,* mitotic cell cycle); protein metabolism; response to lipids; programmed cell death; signal transduction; system development and organizational pathways (*e.g.,* cytoskeleton) (Figure 5C). Cytokine response genes influenced by gravidity (n=20) included *IL1A, IFNB1, F3, CXCL3, CCL20,* and *CXCL2.* Genes in this category were upregulated in multigravidas vs. primigravidas. By comparison, HBC genes that were differentially expressed as a function of gravidity were involved in system development, cell-cell signaling, ion transport, response to lipids and positive regulation of cell proliferation. There was a notable lack of inflammatory response pathways. Individual genes that were differentially expressed between multigravidas and primigravidas, included transcription factors (e*.g.,* W*NT5B)*, ion transporter genes (e*.g., KCNMA1*) and major cytokines (e.g., *CCL13* and *IFNG*). Our analyses of placental macrophages from two distinct origins suggest inherent programming influenced by gravidity status.

## DISCUSSION

We purified MIM and HBC populations from two placental compartments in the setting of uncomplicated term deliveries. RNA sequencing revealed stark differences in the transcriptional signatures of these two cell populations. Both MIMs and HBCs expressed diverse combinations of M1 and M2 subtype markers (Figure 3C); interestingly, HBCs expressed significantly higher levels of both M1 and M2 markers than MIMs. We found distinct transcriptional profiles of MIMs and HBCs with mixed M1/M2 gene expression, lending credence to ongoing discussions that the M1/M2 dichotomy (*33*) does not fully capture the heterogeneous profiles of placental immune cells. Furthermore, much of what is known about macrophage function has been developed through *in vitro* stimulation experiments that may not accurately reflect the *in vivo* microenvironment. Unbiased “-omics” approaches may reveal the broad spectrum of activation states of these cells at the maternal-fetal interface. Overall, our findings highlight the divergent phenotypes of MIMs and HBCs in healthy pregnancies at term, which likely reflects their separate origins and adaptability to changing microenvironments in the distinct compartments they occupy throughout pregnancy.

Most studies to date have suggested that HBCs predominantly exhibit an anti-inflammatory, M2-like phenotype (*15, 28, 40–44*). However, our data demonstrate that HBCs exhibit greater phenotypic heterogeneity than previously described. HBCs expressed both M1 (e.g. *IL6* and *CXCL8*) and M2 (e.g. *IL10* and *VEGF*) markers at high levels (Figure 4C). This is consistent with limited functional analyses of HBCs showing that they express high levels of toll-like receptors and secrete the proinflammatory cytokines IL-6 and IL-8 in response to inflammatory stimuli (*66*). Other studies have demonstrated phagocytic activity of HBCs across all trimesters (*21, 35, 36, 72*). Comparison of cytokine profiles of HBCs from early and late gestation has shown increased expression of inflammatory mediators as pregnancy progresses (*73*). The broad expression of M1 markers in our study also suggests that there may be other pro-inflammatory functional roles for HBCs in normal term placentas. Future studies of HBCs in different disease states, especially leveraging single cell approaches, will help to further elucidate the full spectrum of HBC functional states.

Our results showed that *TWIST1* was 51-fold more highly expressed in MIMs compared to HBCs, which was confirmed by qRT-PCR. Expression of this molecule has been linked to the downregulation of NFKB mediated pro-inflammatory cytokine production in both murine and human macrophages (*74, 75*). *In vitro* studies investigating human monocyte derived macrophages revealed that *TWIST1* downregulated cytokine responses to NOD2, a sensor of bacterial peptidoglycan, through an epigenetic mechanism that enabled the formation of immune memory responses (*76, 77*). Thus, an anti-inflammatory phenotype of MIMs may be regulated by epigenetic modifications. Other differentially expressed regulators (e.g., *NLRP10* and *SLUT2B1*) may be similarly regulated.

Most investigations focusing on MIMs to date been limited to various disease states such as malaria, preeclampsia, and obesity (*24, 30, 49*). We found 13-fold higher expression of *TGM2* and 5-fold higher expression of *SNAP23* in MIMs compared to HBCs*. TGM2* is an anti-inflammatory macrophage marker in both human and mouse (*78*). In mouse studies, *TGM2* plays a protective role in LPS-induced apoptosis of macrophages (*79*). *SNAP23* regulates phagosome formation and maturation in macrophages as well as TLR4 transport upon LPS stimulation (*80, 81*). We also found that MIMs are mixture of M1/M2 subtypes at term (*57, 82*). It is possible that the upregulation of biological processes related to metabolism is indicative of a transition that occurs at parturition (*29*). More studies are needed to fully understand the phenotypic changes that occur during normal pregnancy and birth, as well as in the context of pregnancy complications such as preterm birth.

We also explored the relationship between maternal gravidity and gene expression in MIMs and HBCs (Figure 5). In both cell types, increased gravidity was associated with differential expression of genes related to multiple pathways, including metabolism, development, and cell signaling. However, only five genes were differentially expressed in both MIMs and HBCs as a function of gravidity. Interestingly *MSC* and *C15ORF48* were both more highly expressed in multigravida compared to primigravida; both genes have been linked to suppression of pro-inflammatory response (*83–85*). Our analyses suggest a novel influence of gravidity on the gene signatures of MIMs and HBCs. As many pregnancy complications, such as preeclampsia, are more common in primigravida, we hypothesize that epigenetic mechanisms involved in gene regulation and programming shape the gene signatures and innate memory of MIMs, which may work to facilitate success of subsequent pregnancies. Furthermore, the epigenetic profiles of MIMs and other placental immune cells in multigravid environments could also affect the gene signatures of fetal HBCs. Though multiple studies have delved into the vast epigenetic landscape that shapes normal and disease states in pregnancy (*31, 86, 87*), future work is needed to determine how gravidity influences MIMs and HBCs throughout pregnancy in this context.

Our profiles of maternal-derived MIMs and fetal-derived HBCs are consistent with recent work from Vento-Tormo *et al.* that identified distinct subsets of placental macrophage populations using single-cell RNA sequencing (scRNA-seq) (*56*). Many of the most highly differentially expressed genes we reported for MIMs and HBCs were also identified by their study (**Supplemental** Figure 1). The differences between the markers we report and the scRNA-seq study could be due to differences in the methods of analyses. We directly compared purified MIMs and HBCs vs. the Vento-Tormo *et al*. study that profiled all cells at the maternal-fetal interface, including MIMs and HBCs. Another significant difference was the gestational age of sampling. Vento-Tormo, *et al.* analyzed first trimester vs. term samples, which were the focus of our study.

Thomas *et al.* have recently proposed that earlier studies on HBCs have been contaminated by a population of maternal myeloid cells they termed placental associated maternal macrophages (PAMMs) that localize to the placental surface (*36*). PAMM contamination is unlikely in our study, as our HBC purification technique discarded the first two rounds of tissue digestion containing the syncytial layer and any associated PAMMs. Our methods combined with those described by Thomas *et al*. may simplify identification and isolation of HBCs for future transcriptomic, epigenetic and functional studies.

Our study had limitations. The prevalence of M1 genes and pro-inflammatory markers in MIMs and HBCs could be due to parturition, which is a pro-inflammatory state. Studies suggest that maternal macrophages exhibiting an M1 phenotype promote cervical ripening, uterine contractions, and delivery (*57, 88, 89*). It is also possible that the switch to an M1-subtype helps protect the fetus and mother from infection following amniotic sac rupture and prevents placental retention. Also, studies report an increase in activated monocytes in preterm labor as compared to normal pregnancies (*90*), additional evidence that activated macrophages may play a significant role during parturition. Finally, this study was focused on placentas from African patients with likely lifetime exposures to malaria and many other infectious agents that are endemic to the region. Thus, the results might not be fully representative of those in other populations.

The primary strengths of our study were the isolation and purification of MIMs and HBCs using a targeted cell approach based on placental anatomy and the comparisons of samples from primigravidas and multigravidas. We demonstrated that maternal and fetal myeloid cells at the maternal-fetal interface exhibit diverse gene signatures, with greater transcriptional heterogeneity than previously reported. Our results also suggest a novel influence of maternal gravidity on the transcriptional states of both MIMs and HBCs. Understanding the interactions between these two cell populations is critical to uncover mechanisms of immunoregulation in the context of reproductive memory and placental disease.

## MATERIALS AND METHODS

### Study design

This is a nested case control study of pregnant patients that delivered from January to February 2015 who were enrolled in a randomized controlled trial of intermittent preventative treatment for malaria in pregnancy in Tororo, Uganda (ClinicalTrials.gov number, NCT02163447). Primigravid participants were compared to multigravida participants.

Eligibility criteria included healthy, HIV-negative patients ≥ 16 years of age between 12-20 weeks gestational age at the time of enrollment. Participants were enrolled from June through October 2014. Gestational age was confirmed by ultrasound. Patients were screened for malaria at the time of enrollment and monitored on a monthly basis throughout pregnancy and at delivery. All patients labored and delivered at term. Within 30 minutes of birth, placentas were gently rinsed twice with cold PBS to remove blood clots and debris. Maternal and fetal monocyte/macrophages were isolated as described below. All participants had no evidence of past or active placental malaria, confirmed by placental histopathology (Rogerson criteria) (*91*). Patient characteristics were described in **Supplemental Table 5**.

### Ethics Statement

Written informed consent was obtained from all study participants. Ethical approval was obtained from the Uganda National Council of Science and Technology, the Makerere University School of Medicine Research and Ethics Committee, the Makerere University School of Biomedical Sciences Research and Ethics Committee, and the University of California, San Francisco.

### Isolation of maternal intervillous monocytes (MIMs)

As described previously(*92*) with modifications, maternal blood was collected from the intervillous space by flushing spiral arteries from the maternal surface of the placenta with 30 ml of cold PBS, allowing fluid to passively drain via gravity for 15 minutes. Viable monocytes were isolated in a two-step process: 1) density gradient centrifugation using Ficoll; and 2) depletion of T cells, B cells, NK cells, dendritic cells, granulocytes, and erythrocytes using the Dynabeads Untouched Human Monocytes Kit (Invitrogen, 11350D) (**Supplemental** Figure 2).

### Isolation of placental fetal Hofbauer cells (HBCs)

As previously reported(*35*) with modifications, placentas were washed with cold cytowash to remove excess blood prior to removal of amniotic membranes and decidual tissue. The remaining chorionic villi were minced into 5 mm-sized pieces and washed with cold PBS until clear. At least 50 g of villous tissue was treated with collagenase (0.075% collagenase, 0.04 % DNase, 0.07% hyaluronidase, 3mM CaCl2) for 30 minutes in a 37°C water bath with gentle stirring. Immediately thereafter, the samples were centrifuged (1300 rpm) and the supernatant containing primarily syncytium was discarded. Undigested tissue was resuspended in cold cytowash, and digested with trypsin (0.125% trypsin, 0.02% DNase) for 60 minutes at 37°C, with gentle mixing every 10 minutes. After 60 minutes, additional collagenase was added and the digestion continued for another 30 minutes, also with gentle mixing every 10 minutes. The tissue was disaggregated by pipetting with a 5ml pipette, then filtered over 2x gauze then 1 mm and 90 μM sieves to remove undigested remnants. The cells were collected and washed 4 times with cold cytowash. The cell pellet was applied to a discontinuous Percoll gradient and the interface between 20% and 35% was collected. These cells were washed twice in cytowash, and HBCs were isolated by negative selection on Dynabeads coated with anti-EGFR to remove trophoblasts (Santa Cruz Biotechnology, sc-120) and anti-CD10 to remove fibroblasts (BioLegend, 312202; Goat anti-mouse IgG, Invitrogen, 110-33). The cells were collected, washed in PBS, and stored in RNAlater (Invitrogen) at −80°C (**Supplemental** Figure 1).

### RNA isolation

RNA was extracted from isolated MIMs and HBCs using a RNeasy Micro Kit (Qiagen) following the standard manufacturer’s protocol. RNA quality was assessed using an Agilent 2100 BioAnalyzer (RIN > 7).

### RNA sequencing

We profiled the transcriptomes of six paired samples of MIMs and HBCs by bulk RNA sequencing (RNA-seq) at the UCLA Clinical Microarray Core Facility. cDNA and library construction were conducted using 1 µg RNA per sample and sequenced on an Illumina HiSeq 2500 to obtain ∼30 million reads per sample.

### Quantitative RT-PCR

We investigated expression levels of target genes to validate our profiling results and further explore relationships across a larger set of samples. In total, we examined 19 MIM and 15 HBC samples (**Supplemental Table 5)**. These were a mixture of remaining samples from our RNA-seq analysis (MIMs, n=6; HBCs, n=2;) and additional samples from independent placentas collected concurrently (MIMs, n=13; HBCs, n=13;). We converted purified RNA samples to cDNA using iSCRIPT Universal TaqMan (Bio-Rad), and performed qRT-PCR using TaqMan primers for selected targets (**Supplemental Table 6**) mixed with TaqMan Universal Master Mix II, no UNG (Life Technologies, Quant Studio 6). Reactions were carried out for 40 cycles. At least 3 technical replicates were analyzed for all comparisons. Differential expression between HBCs vs. MIMs was calculated via the ΔΔCT method. We normalized expression using the mean CT of housekeeping genes, *GAPDH* and *ACTB*.

### Statistical Analysis

For RNAsequencing, FASTQ files were aligned to the human reference genome (GRCh37/hg19) using BWA (Burrows-Wheeler Alignment tool). Aligned BAM files were processed using htseqcount to obtain counts-per-million (CPM) values. Genes with CPM values > 0.5 in at least two samples were considered expressed above background. Data were further transformed using VOOM (*93*) and differentially expressed transcripts were identified using LIMMA. Genes missing a HGNC symbol were excluded in downstream analyses. In total, we examined 18,802 unique RNAs using this approach. We identified differentially expressed transcripts based on: 1) cell type of origin (MIMs vs. HBCs); or 2) cell type vs. gravidity. We defined differentially expressed transcripts between MIMs and HBCs as p<0.00001 (unadjusted); absolute FC ≥ 4, false discovery rate <0.1%). In gravidity-based analyses, we considered genes expressed with high confidence in each cell type (average CPM > 0.5) and applied a cutoff of unadjusted p<0.05, absolute FC ≥ 2. We conducted principal components analysis and hierarchical clustering of FC values using average linkage and Euclidean distance (pheatmap, RRID:SCR_016418) (*94*). Cell/tissue enrichment of the most abundant top 2% genes in each cell type was conducted using CTen (*95*). Functional enrichment analysis of Gene Ontology (GO) Biological Processes was evaluated via DAVID and a cutoff criteria of p<0.01; fold enrichment > 1.5, and minimum differentially expressed transcripts ≥10 (MIMs vs. HBCs) or ≥5 (cell type vs. gravidity) per GO term. Terms were grouped based on GO classifications. Raw and normalized data were deposited in the NCBI Gene Expression Omnibus **(GEO; TBD).**

For qRT-PCR, we employed Dunnett’s test (SSPS) to determine significant differences in expression (p<0.05). Relative FC values were expressed as average log2 ratios between MIMs and HBCs.

### List of Supplementary Materials

**1- Figure S1. Hierarchical clustering of Hofbauer cell markers defined by Vento-Tormo et. al. (A)** Out of 30 markers identified in the single cell RNA-seq study of Vento-Tormo et.al, 19 were found to be significant according to our analysis (adj-p<0.01). Fold changes for each sample were calculated according to average MIM value. **(B)** Using the significant 19 markers, a new fold change between HBCs-MIMs was calculated for each sample pair.
**2- Figure S2. Experimental approach to isolate macrophages in human term placentas.** Workflow applied to acquire fetal and maternal macrophages from human placentas for transcriptomic analyses. In brief, term placentas were collected from Tororo, Uganda. MIMs were isolated from intervillous blood using Ficoll centrifugation and negative selection with magnetic beads. HBCs were obtained through trypsin/collagenase enzymatic digestion of dissected villous tissues, Percoll density centrifugation, and negative section with magnetic beads (CD10, EGFR). RNA-sequencing was performed using purified RNA from MIM and HBC samples.
**3- Table S1. Relative gene expression differences between MIMs and HBCs via RNA-seq and qRT-PCR.** Comparisons of fold change (FC) and significance of differentially expressed genes between MIMs and HBCs identified via RNA-seq and qRT-PCR Pearson’s correlation between the two datasets was 0.97. Genes shaded in grey were not defined as significant based on separate criteria for the two approaches: qRT-PCR, p<0.05; RNA-seq, p<0.00001; absolute log2FC > 2.
**4- Table S2. Gene ontology analysis of differentially expressed genes between MIMs and HBCs.** Significant biological processes identified via DAVID (p<0.01; number changed >10). Number of differentially expressed genes higher in MIMs or HBCs are noted. Ratio between the two subsets. P-value regarding significance and list of genes differentially expressed in MIMs or HBCs.
**5- Table S3. Differential expression of M1/M2 markers in MIMs vs. HBCs.** A panel of 19 anti-inflammatory (M1) and 21 pro-inflammatory (M2) markers were identified based on a broad literature review. Out of this panel, 24 genes were observed to be differentially expressed between MIMs and HBCs (average expression>0, p<0.01 and absolute log2FC > 1; please note less conservative criteria than unbiased analysis). Abbreviations include: Log2 average counts in MIMs (MIM_Ct) or HBCs (HBC_Ct); fold change difference between MIMs and HBCs (FC, log2); and significance (p-value). Red color indicates significance.
**6- Table S4. Gene ontology analysis of genes Influenced by gravidity in MIMs or HBCs.** Significant biological processes influenced by gravid status in MIMs or HBCs identified via DAVID. Criteria: p<0.01 in either MIMs or HBCs and minimum differentially expressed genes ≥ 10.
**7- Table S5. Patient characteristics and birth outcomes and experimental usage.**
**8- Table S6. List of Taqman qRTPCR probes used for validation studies.**

## ACKNOWLEDGMENTS

We thank the patients who participated in this study. We also thank John Ategeka and Samuel Wamala for technical support.

## Funding

National Institutes of Allergy and Infectious Diseases grant K08AI141728 (SLG)

National Institutes of Allergy and Infectious Diseases grant K24AI113002 (MEF)

Eunice Kennedy Shriver National Institute of Child Health and Human Development

Reproductive Scientist Development Program grant K12 HD000849 (SLG)

Eunice Kennedy Shriver National Institute of Child Health & Human Development grant P01HD059454 (GD)

Burroughs Wellcome Fund for the Reproductive Scientist Development Program (SLG)

## Author contributions

Conceptualization: MEF, GC, SJF, SLG

Methodology: JFR, GC, PJR, MEF, SJF, SLG

Investigation: NO, JFR, SB, MYK, MRA, JA, DJM, SLG

Visualization: NO, JFR, SB, MRA, JA

Funding acquisition: MRK, GD, GC, MEF, SJF, SLG

Project administration: MRK, AK, GD, MEF, SLG

Supervision: MRK, AK, GD, PJR, MEF, SLG

Writing – original draft: NO, JFR, MRA, SLG

Writing – review & editing: NO, JFR, SB, MYK, MRA, JA, DJM, MRK, AK, GD, PJR, GC, MEF, SJF, SLG

## Data and materials availability

RNA-seq data has been uploaded on Gene Expression Omnibus (waiting for accession number).

