## Supplemental Figures and Tables for "Distinct transcriptional profiles of maternal and fetal placental macrophages at term are associated with gravidity"

### Supplemental Figure 1.

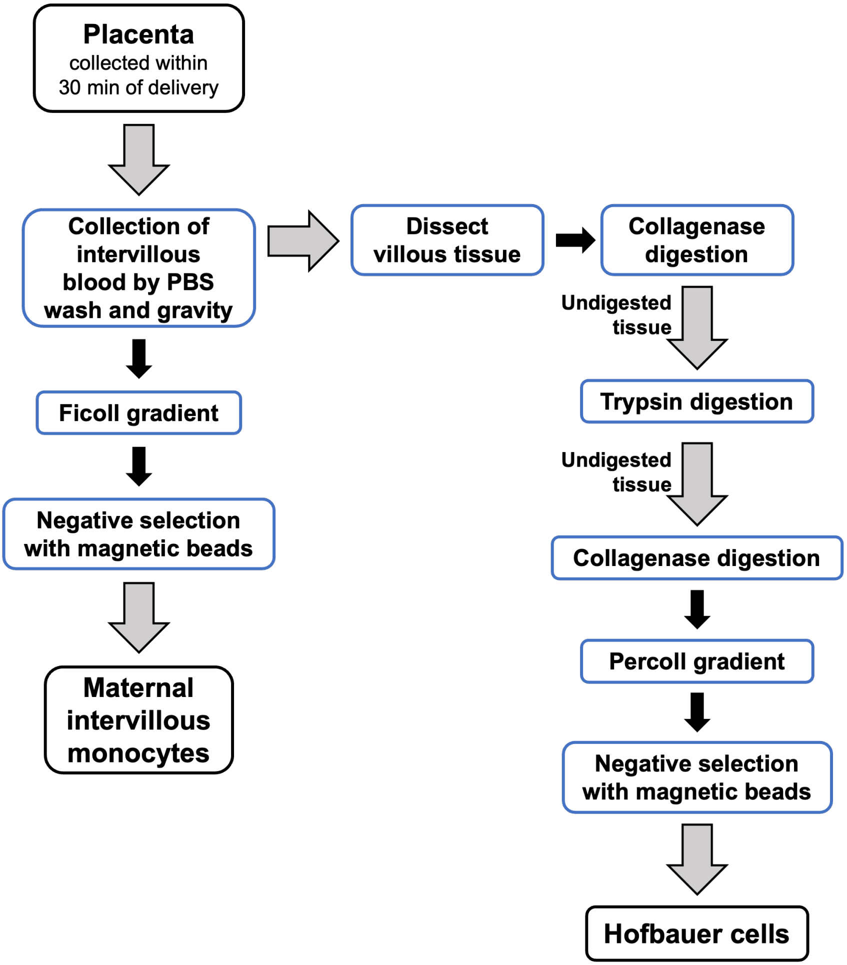

**Supplemental Figure 2. Hierarchical clustering of Hofbauer cell markers defined by Vento-Tormo et. al.**
(**A**) Out of 30 markers identified in the single cell RNA-seq study of Vento-Tormo et.al, 19 were found to be significant according to our analysis (adj-p<0.01). Fold changes for each sample were calculated according to average MIM value. (**B**) Using the significant 19 markers, a new fold change between HBCs-MIMs was calculated for each sample pair.

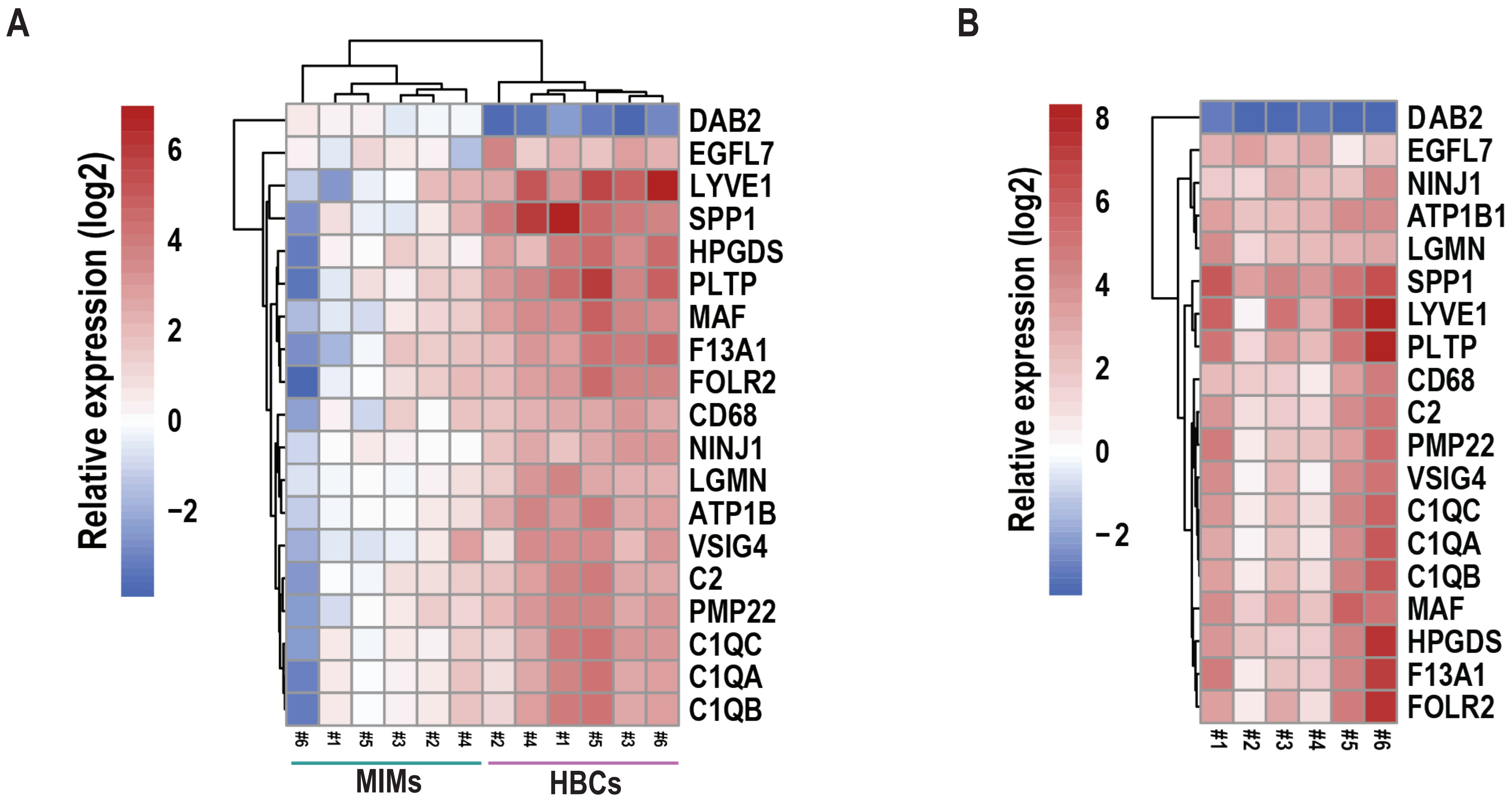

**Supplemental Table 1. Patient characteristics and birth outcomes and experimental usage.**

| Patient ID | Maternal age (years) | Gestational age (weeks) | Gravidity | Mode of Delivery | Birth weight (gram) | Low birth weight (<2500 g) | Small for gestational age | Infant Sex | RNA-seq experiment | qRT-PCR experiment | |
|---|---|---|---|---|---|---|---|---|---|---|---|
| | | | | | | | | | | MIMs | HBCs |
| #1 | 22 | 41.0 | Multigravida | Vaginal | 3450 | No | No | Male | Yes | Yes | No |
| #2 | 18 | 39.9 | Primigravida | Vaginal | 2950 | No | No | Female | Yes | Yes | No |
| #3 | 27 | 40.6 | Multigravida | Vaginal | 3580 | No | No | Female | Yes | Yes | No |
| #4 | 18 | 41.3 | Primigravida | Cesarean | 3140 | No | No | Male | Yes | Yes | Yes |
| #5 | 26 | 40.6 | Multigravida | Vaginal | 2830 | No | No | Male | Yes | Yes | Yes |
| #6 | 18 | 37.1 | Primigravida | Vaginal | 2670 | No | No | Male | Yes | Yes | No |
| #7 | 20 | 40.6 | Primigravida | Cesarean | 2800 | No | No | Male | No | No | Yes |
| #8 | 29 | 38.0 | Multigravida | Vaginal | 2230 | Yes | Yes | Male | No | Yes | Yes |
| #9 | 22 | 41.1 | Multigravida | Vaginal | 2620 | No | Yes | Female | No | Yes | Yes |
| #10 | 29 | 39.0 | Multigravida | Vaginal | 2520 | No | Yes | Male | No | Yes | Yes |
| #11 | 22 | 39.4 | Multigravida | Vaginal | 3400 | No | No | Male | No | Yes | Yes |
| #12 | 25 | 39.0 | Multigravida | Vaginal | 2780 | No | No | Male | No | Yes | No |
| #13 | 22 | 40.3 | Multigravida | Vaginal | 2850 | No | No | Female | No | Yes | Yes |
| #14 | 19 | 41.1 | Multigravida | Cesarean | 2850 | No | Yes | Female | No | Yes | Yes |
| #15 | 26 | 40.9 | Multigravida | Vaginal | 3090 | No | No | Male | No | Yes | Yes |
| #16 | 20 | 37.7 | Multigravida | Vaginal | 2880 | No | No | Female | No | Yes | Yes |
| #17 | 28 | 39.0 | Multigravida | Vaginal | 3029 | No | No | Male | No | Yes | No |
| #18 | 31 | 39.0 | Multigravida | Vaginal | 3420 | No | No | Male | No | Yes | Yes |
| #19 | 27 | 43.0 | Multigravida | Vaginal | 3120 | No | No | Male | No | Yes | Yes |
| #20 | 25 | 39.0 | Multigravida | Vaginal | 3150 | No | No | Male | No | Yes | Yes |
| #21 | 22 | 39.4 | Multigravida | Vaginal | 2900 | No | No | Male | No | No | Yes |

**Supplemental Table 2. List of Taqman qRT-PCR probes used for validation studies.**

| Gene | Taqman identifier |
|---|---|
| GREM1 | Hs01879841_s1 |
| TWIST1 | Hs00361186_m1 |
| CYP19A1 | Hs00903413_m1 |
| CCL2 | Hs00234140_m1 |
| S1PR1 | Hs00173499_m1 |
| IL6 | Hs00985639_m1 |

**Supplemental Table 3. Relative gene expression differences between MIMs and HBCs via RNA-seq and qRT-PCR.** Comparisons of fold change (FC) and significance of differentially expressed genes between MIMs and HBCs identified via RNA-seq and qRT-PCR Pearson's correlation between the two datasets was 0.97. Genes shaded in grey were not defined as significant based on separate criteria for the two approaches: qRT-PCR, $p<0.05$; RNA-seq, $p<0.00001$; absolute $\log_2$FC > 2.

| **HBCs vs. MIMs** | **TWIST1** | **CYP19A1** | **CCL2** | **S1PR1** | **IL6** | **GREM1** |
|---|---:|---:|---:|---:|---:|---:|
| $Log_2$FC (qRT-PCR) | -4.87991 | -4.12415 | 1.389308 | 1.10748 | 1.902728 | 3.193769 |
| p-value (qRT-PCR) | 9.82E-12 | 8.6E-08 | 0.000145 | 0.003528 | 0.00048 | 2.23E-07 |
| $Log_2$FC (RNA-seq) | -5.67689 | -4.29884 | 3.58 | 4.46 | 3.492038 | 3.863052 |
| p-value (RNA-seq) | 9.5E-12 | 8.32E-07 | 4.47E-06 | 5.76E-07 | 0.002403 | 0.003953 |

**Supplemental Table 4: Gene ontology analysis of differentially expressed genes between MIMs and HBCs.** Significant biological processes identified via DAVID (p<0.01; number changed >10). Number of differentially expressed genes higher in MIMs or HBCs are noted. Ratio between the two subsets. P-value regarding significance and list of genes differentially expressed in MIMs or HBCs.

| Term | MIM | HBC | Ratio | ALL (p) | MIM_up | HBC_up |
|---|---|---|---|---|---|---|
| cell motility | 50 | 49 | 1.0 | 5.4E-07 | CGA, PRKCZ, PPP2R3A, PGF, TBX20, WWC1, SLC7A8, GIPC1, POSTN, RDX, CCL28, PTEN, STARD13, PFN2, DAB2, P2RY6, ZNF703, PAK3, CCSAP, GAB1, SEMA3B, NOS3, RAB25, DEPDC1B, RHOD, DCX, NET1, CYP19A1, TWIST1, PARD6B, LURAP1, VAV3, S100P, ARTN, MIEN1, HES1, SDC1, SVBP, RRAS2, OPHN1, HSPB1, WASL, APBB2, ADGRL3, GRB7, EMP2, TNFAIP1, NLRP10, GSTP1, KALRN | NRP2, ADCY3, ATP1B1, JAG2, JAG1, SLC7A5, IL10, SLC16A1, S1PR1, UNC5C, RAPGEF2, IL1A, DPP4, MATK, LDB2, PDE4D, LDLRAD4, DDIT4, TNFAIP6, MMP10, CCR7, CCR4, SEMA4C, SRGAP3, NKD1, CCL2, SPOCK2, CXCL3, CSF1, KIT, SRC, HRH1, IL23A, CCL20, ITGAV, TNFRSF18, CD2, ZC3H12A, HSPA5, NFATC2, OLR1, SPHK1, SMAD3, EVL, ADGRG1, PLCG1, ITGA6, ITGA5, ABL2 |
| cell migration | 46 | 41 | 0.9 | 5.1E-06 | CGA, PRKCZ, PGF, TBX20, WWC1, SLC7A8, GIPC1, RDX, POSTN, PTEN, CCL28, STARD13, P2RY6, DAB2, PFN2, ZNF703, PAK3, SEMA3B, NOS3, RAB25, DEPDC1B, RHOD, DCX, NET1, TWIST1, CYP19A1, LURAP1, S100P, VAV3, ARTN, MIEN1, HES1, SDC1, SVBP, RRAS2, OPHN1, HSPB1, WASL, APBB2, EMP2, ADGRL3, GRB7, GSTP1, NLRP10, TNFAIP1, KALRN | NRP2, ATP1B1, CCL2, CXCL3, CSF1, KIT, SLC7A5, IL10, SRC, HRH1, SLC16A1, S1PR1, IL23A, CCL20, ITGAV, TNFRSF18, CD2, ZC3H12A, HSPA5, NFATC2, RAPGEF2, DPP4, IL1A, MATK, OLR1, SPHK1, SMAD3, PDE4D, EVL, ADGRG1, LDLRAD4, DDIT4, TNFAIP6, CCR7, ITGA6, PLCG1, ITGA5, CCR4, SEMA4C, SRGAP3, ABL2 |
| regulation of cell motility | 29 | 35 | 1.2 | 5.6E-07 | CGA, PPP2R3A, PGF, POSTN, RDX, PTEN, CCL28, STARD13, DAB2, PFN2, P2RY6, ZNF703, PAK3, CCSAP, GAB1, SEMA3B, RAB25, RHOD, CYP19A1, TWIST1, PARD6B, MIEN1, SVBP, RRAS2, HSPB1, WASL, GRB7, EMP2, GSTP1 | NRP2, NKD1, CCL2, SPOCK2, CXCL3, CSF1, JAG2, JAG1, KIT, SRC, S1PR1, IL23A, CCL20, ITGAV, TNFRSF18, ZC3H12A, HSPA5, UNC5C, RAPGEF2, IL1A, SPHK1, LDB2, SMAD3, EVL, ADGRG1, LDLRAD4, MMP10, TNFAIP6, CCR7, ITGA6, PLCG1, ITGA5, SEMA4C, SRGAP3, ABL2 |
| regulation of cell migration | 26 | 32 | 1.2 | 4.8E-06 | CGA, PGF, POSTN, RDX, PTEN, CCL28, STARD13, PFN2, DAB2, P2RY6, ZNF703, PAK3, GAB1, SEMA3B, RAB25, RHOD, CYP19A1, PARD6B, MIEN1, SVBP, RRAS2, HSPB1, WASL, GRB7, EMP2, GSTP1 | NRP2, CCL2, CXCL3, CSF1, JAG2, JAG1, KIT, SRC, S1PR1, IL23A, CCL20, ITGAV, TNFRSF18, ZC3H12A, HSPA5, UNC5C, RAPGEF2, IL1A, SPHK1, LDB2, SMAD3, EVL, ADGRG1, LDLRAD4, TNFAIP6, MMP10, CCR7, ITGA6, PLCG1, ITGA5, SEMA4C, SRGAP3 |
| chemotaxis | 14 | 27 | 1.9 | 2.0E-03 | VAV3, STX3, ANK3, PGF, ARTN, OPHN1, HSPB1, SIAH1, SEMA3B, APBB2, CCL28, ETV4, GSTP1, CYP19A1 | NRP2, CCL2, CXCL3, CSF1, KIT, IL10, HRH1, S1PR1, IL23A, CCL20, BCL11B, ITGAV, CXCR6, UNC5C, DRAXIN, SMAD3, EVL, PDE4D, NCAM1, CCR8, CCR7, CCR4, NTRK1, SEMA4C, SPTBN1, SPTAN1, FEZ1 |
| (+) regulation of cell motility | 16 | 24 | 1.5 | 7.4E-06 | CGA, PGF, POSTN, RDX, MIEN1, P2RY6, DAB2, ZNF703, PAK3, RRAS2, HSPB1, SEMA3B, RAB25, RHOD, GRB7, TWIST1 | NRP2, CCL2, SPOCK2, CSF1, CXCL3, SPHK1, SMAD3, KIT, SRC, TNFAIP6, CCR7, S1PR1, IL23A, ITGA6, PLCG1, CCL20, ITGA5, ITGAV, SEMA4C, TNFRSF18, ZC3H12A, HSPA5, RAPGEF2, IL1A |
| (+) regulation of cellular component movement | 16 | 24 | 1.5 | 1.4E-05 | CGA, PGF, POSTN, RDX, MIEN1, P2RY6, DAB2, ZNF703, PAK3, RRAS2, HSPB1, SEMA3B, RAB25, RHOD, GRB7, TWIST1 | NRP2, CCL2, SPOCK2, CSF1, CXCL3, SPHK1, SMAD3, KIT, SRC, TNFAIP6, CCR7, S1PR1, IL23A, ITGA6, PLCG1, CCL20, ITGA5, ITGAV, SEMA4C, TNFRSF18, ZC3H12A, HSPA5, RAPGEF2, IL1A |
| (+) regulation of cell migration | 15 | 23 | 1.5 | 1.9E-05 | CGA, PGF, POSTN, RDX, MIEN1, DAB2, P2RY6, ZNF703, PAK3, RRAS2, HSPB1, RAB25, SEMA3B, RHOD, GRB7 | NRP2, CCL2, CSF1, CXCL3, SPHK1, SMAD3, KIT, SRC, TNFAIP6, CCR7, S1PR1, IL23A, ITGA6, PLCG1, CCL20, ITGA5, ITGAV, SEMA4C, TNFRSF18, ZC3H12A, HSPA5, RAPGEF2, IL1A |
| single organismal cell-cell adhesion | 20 | 32 | 1.6 | 9.6E-04 | PRKCZ, GLDN, NECTIN3, CAMSAP3, RDX, BAD, PAWR, CSRP1, SOD1, CCL28, HES1, IGSF5, ZNF703, PAK3, ANK3, PDE5A, TTYH1, HSPB1, TSTA3, DLG5 | CCL2, CD8A, IL6ST, JAG2, KIT, IL7R, TNFRSF4, IL10, SRC, IL23A, ICOS, BCL11B, ITGAV, MAP3K8, CD2, CD6, CD5, DPP4, CD28, CD7, FZD8, ITK, TCF7, CD3E, CTLA4, SMAD3, NLGN2, CCR7, ITGA6, ITGA5, ITGAD, PRNP |

| | | | | | | |
|---|---|---|---|---|---|---|
| (+) regulation of cell adhesion | 13 | 24 | 1.8 | 2.6E-05 | PRKCZ, STX3, VAV3, CYTH3, BAD, CCL28, HES1, ANK3, PAK3, TGM2, RHOD, EMP2, NET1 | CCL2, IL6ST, SPOCK2, CD3E, CSF1, CTLA4, SMAD3, NINJ1, IL7R, ADGRG1, IL10, SRC, CCR7, IL23A, ITGA6, ITGA5, ITGAV, ICOS, MAP3K8, TNFRSF18, CD6, CD5, DPP4, CD28 |
| regulation of cell-cell adhesion | 11 | 18 | 1.6 | 8.5E-03 | HES1, PRKCZ, ZNF703, PAK3, ANK3, PDE5A, RDX, BAD, PAWR, SOD1, CCL28 | CCL2, CD3E, IL6ST, CTLA4, IL7R, IL10, SRC, CCR7, IL23A, ITGA6, ICOS, MAP3K8, CD2, PRNP, CD6, CD5, DPP4, CD28 |
| (+) regulation of cell-cell adhesion | 5 | 16 | 3.2 | 8.2E-03 | HES1, PRKCZ, PAK3, ANK3, BAD | CCL2, CD3E, IL6ST, CTLA4, IL7R, IL10, SRC, CCR7, IL23A, ITGA6, ICOS, MAP3K8, CD6, CD5, DPP4, CD28 |
| cell-matrix adhesion | 9 | 11 | 1.2 | 1.0E-03 | PRKCZ, SORBS1, LYPD5, POSTN, RHOD, EMP2, PTEN, CCL28, RASA1 | CD96, CCR7, LYPD3, ITGA6, EPDR1, ITGB8, ITGAV, CSF1, SMAD3, NINJ1, SRC |
| regulation of cell-matrix adhesion | 7 | 5 | 0.7 | 2.9E-03 | PRKCZ, POSTN, RHOD, EMP2, PTEN, CCL28, RASA1 | CCR7, CSF1, SMAD3, NINJ1, SRC |
| inflammatory response | 8 | 38 | 4.8 | 1.7E-03 | PRKCZ, SDC1, VAMP8, TGM2, SNAP23, CDO1, GSTP1, CYP19A1 | CCL2, LDLR, IL6ST, TNFRSF25, CXCL3, CSF1, DUSP10, NFKB1, AFAP1L2, NFKB2, KIT, TNFRSF4, IL10, CD96, HRH1, IL23A, REL, CCL20, PTGES, TICAM1, CXCR6, TNFRSF18, ZC3H12A, CD6, IL1A, CD28, TNIP3, IRAK2, OLR1, IL1RL1, SPHK1, SMAD3, TNFRSF9, TNFAIP6, CCR7, CCR4, KDM6B, IGFBP4 |
| leukocyte cell-cell adhesion | 9 | 28 | 3.1 | 2.6E-03 | HES1, PRKCZ, PAK3, PDE5A, TSTA3, BAD, PAWR, SOD1, CCL28 | CCL2, CD8A, IL6ST, JAG2, KIT, IL7R, TNFRSF4, IL10, SRC, IL23A, ICOS, BCL11B, MAP3K8, CD2, CD6, CD5, DPP4, CD7, CD28, FZD8, ITK, TCF7, CD3E, CTLA4, SMAD3, CCR7, ITGA5, PRNP |
| T cell activation | 7 | 27 | 3.9 | 3.4E-03 | HES1, PRKCZ, PAK3, PDE5A, BAD, PAWR, SOD1 | CCL2, CD8A, IL6ST, JAG2, KIT, IL7R, TNFRSF4, IL10, SRC, IL23A, ICOS, BCL11B, MAP3K8, CD2, CD6, CD5, DPP4, CD7, CD28, FZD8, ITK, TCF7, CD3E, CTLA4, SMAD3, CCR7, PRNP |
| leukocyte migration | 9 | 23 | 2.6 | 8.2E-04 | VAV3, PGF, SLC7A8, ARTN, WASL, GRB7, CCL28, NLRP10, CYP19A1 | ATP1B1, CCL2, OLR1, CSF1, CXCL3, PDE4D, KIT, SLC7A5, IL10, SRC, CCR7, SLC16A1, HRH1, S1PR1, IL23A, ITGA6, PLCG1, CCL20, ITGA5, ITGAV, TNFRSF18, CD2, IL1A |
| (+) regulation of cytokine production | 7 | 24 | 3.4 | 2.5E-03 | PRKCZ, LURAP1, HSPB1, POSTN, SOD1, NLRP10, TWIST1 | IRAK1, CCL2, PANX1, IL6ST, CD3E, IL1RL1, SMAD3, NFKB1, AFAP1L2, PDE4D, NFKB2, IL10, SRC, CCR7, IL23A, CCL20, TICAM1, CD2, HEG1, SPTBN1, CD6, EIF2AK3, IL1A, CD28 |
| response to lipopolysaccharide | 2 | 26 | 13.0 | 5.2E-04 | NOS3, GSTP1 | CSF3, CCL2, TNFRSF25, CXCL3, DUSP10, NFKB1, ABCA1, NFKB2, TNFRSF4, IL10, NOCT, SRC, CD96, CCL20, PTGES, TICAM1, TNFRSF18, ZC3H12A, CD6, TNIP3, IRAK2, IRAK1, PDE4D, TNFRSF9, CCR7, PENK |
| response to molecule of bacterial origin | 2 | 26 | 13.0 | 1.0E-03 | NOS3, GSTP1 | CSF3, CCL2, TNFRSF25, CXCL3, DUSP10, NFKB1, ABCA1, NFKB2, TNFRSF4, IL10, NOCT, SRC, CD96, CCL20, PTGES, TICAM1, TNFRSF18, ZC3H12A, CD6, TNIP3, IRAK2, IRAK1, PDE4D, TNFRSF9, CCR7, PENK |
| (+) regulation of leukocyte activation | 6 | 21 | 3.5 | 1.7E-03 | HES1, PRKCZ, VAV3, VAMP8, PAK3, BAD | CCL2, CD3E, IL1RL1, IL6ST, CTLA4, NECTIN2, IL7R, TNFRSF4, IL10, SRC, CCR7, IL23A, ICOS, TICAM1, MAP3K8, CD2, CD6, NFATC2, CD5, DPP4, CD28 |
| (+) regulation of cell activation | 6 | 21 | 3.5 | 2.5E-03 | HES1, PRKCZ, VAV3, VAMP8, PAK3, BAD | CCL2, CD3E, IL1RL1, IL6ST, CTLA4, NECTIN2, IL7R, TNFRSF4, IL10, SRC, CCR7, IL23A, ICOS, TICAM1, MAP3K8, CD2, CD6, NFATC2, CD5, DPP4, CD28 |
| regulation of leukocyte cell-cell adhesion | 8 | 17 | 2.1 | 7.5E-03 | HES1, PRKCZ, PAK3, PDE5A, BAD, PAWR, SOD1, CCL28 | CCL2, CD3E, IL6ST, CTLA4, IL7R, IL10, SRC, CCR7, IL23A, ICOS, MAP3K8, CD2, PRNP, CD6, CD5, DPP4, CD28 |

| | | | | | | |
|---|---|---|---|---|---|---|
| cytokine secretion | 6 | 11 | 1.8 | 3.0E-03 | PRKCZ, POSTN, ANXA4, NLRP10, IL36RN, TWIST1 | TNFRSF9, CCR7, PANX1, IL1RL1, CD2, ZC3H12A, SPTBN1, ABCA1, IL10, SRC, IL1A |
| regulation of cytokine secretion | 6 | 10 | 1.7 | 3.0E-03 | PRKCZ, POSTN, ANXA4, NLRP10, IL36RN, TWIST1 | TNFRSF9, CCR7, PANX1, IL1RL1, CD2, ZC3H12A, SPTBN1, IL10, SRC, IL1A |
| cellular response to molecule of bacterial origin | 2 | 14 | 7.0 | 4.1E-03 | NOS3, GSTP1 | IRAK2, CSF3, IRAK1, CCL2, PDE4D, NFKB1, ABCA1, IL10, SRC, CCL20, TICAM1, ZC3H12A, CD6, TNIP3 |
| interleukin-1 production | 2 | 8 | 4.0 | 4.7E-03 | HSPB1, GSTP1 | CCR7, PANX1, CCL20, SPHK1, SMAD3, ZC3H12A, ABCA1, IL10 |
| programmed cell death | 55 | 53 | 1.0 | 3.2E-03 | HTATIP2, MAEL, PAWR, PTEN, DAB2, AES, DYNLL1, PAK3, NOS3, DLG5, NET1, TWIST1, KLLN, SOCS2, TFPT, CRYAB, LGALS13, CECR2, ARHGEF12, STK3, MIEN1, NME6, RFK, LGALS16, SCIN, GADD45G, LGALS14, TFAP2A, HSPB1, SIAH1, GSTP1, TNFAIP1, KALRN, PRKCZ, GULP1, BEX2, UBE2V2, FIS1, AKT1S1, TGM2, PHLDA3, RASA1, TMEM79, VAV3, TBX3, SMAD6, PDK4, RYBP, BAD, SOD1, ANXA4, AKTIP, CFDP1, GRK5, APBB2 | STIL, IL6ST, LGMN, TNFSF15, JAG2, NFKB1, IL10, TICAM1, MAP3K8, UNC5C, RAPGEF2, IL1A, IRAK1, CD3E, GZMA, DRAXIN, DDIT4, SERPINB9, CTSL, AMIGO2, TNFRSF9, CCR7, CCND2, ERN1, PRNP, EIF2AK3, TRAF1, CCL2, TNFRSF25, HK2, KIT, TNFRSF4, SRC, ITGAV, BCL11B, TNFRSF18, CD2, ZC3H12A, HSPA5, CD5, PHLDA1, CD28, PDK1, IL2RB, TCF7, TP53BP2, SPHK1, CTLA4, SMAD3, ITGA6, ITGA5, NTRK1, DRAM1 |
| regulation of programmed cell death | 45 | 40 | 0.9 | 4.4E-03 | PRKCZ, HTATIP2, MAEL, BEX2, UBE2V2, PAWR, PTEN, FIS1, DAB2, AKT1S1, AES, DYNLL1, PAK3, TGM2, NOS3, DLG5, PHLDA3, RASA1, NET1, TWIST1, VAV3, SOCS2, TBX3, CRYAB, SMAD6, PDK4, LGALS13, BAD, ARHGEF12, SOD1, ANXA4, MIEN1, STK3, LGALS16, SCIN, LGALS14, GADD45G, TFAP2A, HSPB1, SIAH1, CFDP1, GRK5, APBB2, GSTP1, KALRN | TRAF1, STIL, CCL2, IL6ST, TNFRSF25, LGMN, TNFSF15, NFKB1, KIT, TNFRSF4, IL10, SRC, BCL11B, ITGAV, TNFRSF18, ZC3H12A, HSPA5, UNC5C, RAPGEF2, IL1A, IRAK1, IL2RB, TCF7, TP53BP2, CD3E, GZMA, DRAXIN, SPHK1, CTLA4, SMAD3, SERPINB9, AMIGO2, TNFRSF9, CCR7, ITGA6, CCND2, ITGA5, NTRK1, PRNP, EIF2AK3 |
| (-) regulation of cell death | 27 | 31 | 1.1 | 4.8E-03 | PRKCZ, HTATIP2, MAEL, UBE2V2, PTEN, NPAS2, DAB2, TGM2, NOS3, RASA1, TWIST1, TBX3, SOCS2, CRYAB, SMAD6, PDK4, BAD, SOD1, ANXA4, MIEN1, BTBD10, HSPB1, TFAP2A, CFDP1, GRK5, APBB2, GSTP1 | CSF3, STIL, CCL2, IL6ST, LGMN, NFKB1, KIT, IL10, SRC, REL, BCL11B, ITGAV, TNFRSF18, ZC3H12A, HSPA5, IL1A, IRAK1, TCF7, IL2RB, DRAXIN, SPHK1, SMAD3, AMIGO2, SERPINB9, NPC1, CCR7, ITGA6, CCND2, ITGA5, NTRK1, PRNP |
| (-) regulation of programmed cell death | 25 | 28 | 1.1 | 8.6E-03 | PRKCZ, HTATIP2, MAEL, UBE2V2, PTEN, DAB2, TGM2, NOS3, RASA1, TWIST1, TBX3, SOCS2, CRYAB, SMAD6, PDK4, BAD, SOD1, ANXA4, MIEN1, HSPB1, TFAP2A, CFDP1, GRK5, APBB2, GSTP1 | STIL, CCL2, IL6ST, LGMN, NFKB1, KIT, IL10, SRC, ITGAV, BCL11B, TNFRSF18, ZC3H12A, HSPA5, IL1A, IRAK1, TCF7, IL2RB, DRAXIN, SPHK1, SMAD3, AMIGO2, SERPINB9, CCR7, ITGA6, CCND2, ITGA5, NTRK1, PRNP |
| regulation of signal transduction | 86 | 82 | 1.0 | 7.6E-06 | SPIN1, TBX20, WWC1, PIP5K1B, POSTN, ARHGAP17, PAWR, PTEN, DAB2, AES, DYNLL1, IFT20, PAK3, GAB1, NOS3, DEPDC1B, RHOD, RNF146, NET1, RS1, TWIST1, PID1, SOCS2, TRIM40, ARHGEF12, STX1B, STK3, CNGA1, MAP4K3, HES1, ATP6V1C2, DACT2, GADD45G, PDE5A, OPHN1, HSPB1, SIAH1, ADAMTS3, EMP2, TNFAIP1, EPS8L1, GSTP1, KALRN, PRKCZ, EID2, PPP2R3A, RAP1GAP, GLIS2, GIPC1, STARD10, RDX, CYTH3, FAM13A, STARD13, RBX1, PSMB5, MTM1, FIS1, AKT1S1, ZNF703, AMER2, SORBS1, SMARCB1, ARHGAP42, TGM2, RHOBTB1, PHLDA3, RASA1, MTMR4, RNF14, FBXO8, DVL3, LURAP1, VAV3, ERH, SMAD6, BAD, CBY1, TAX1BP3, SOD1, TRIM62, CISH, IL36RN, GRK5, GRB7, PLEKHA1 | NAF1, CD8A, IL6ST, LGMN, EZH2, JAG2, TNFSF15, NFKB1, JAG1, SHE, IL10, SPRY1, SH2D1A, MAP3K8, TICAM1, RAPGEF2, RAMP1, IL1A, TNIP3, IRAK2, IRAK1, TNIK, CD3E, TRABD2A, DRAXIN, PDE4D, PIK3IP1, LDLRAD4, TRAT1, DDIT4, ACVR2A, CCR7, ZFYVE28, SRGAP3, ERN1, SEMA4C, PRNP, PPP1R15B, EIF2AK3, RASD2, PMEPA1, CSF3, TRAF1, NKD1, CCL2, TNFRSF25, CSF1, DUSP10, AFAP1L2, ABCA1, KIT, SESN2, TNFRSF4, SRC, LIF, TSPYL2, IL23A, CCL20, REL, ITGAV, TNFRSF18, ZC3H12A, HSPA5, AXIN2, CD28, FZD8, TP53BP2, IL1RL1, ASXL1, SPHK1, SMAD3, NLGN2, IGSF9B, RGS16, ADGRG1, RGS13, DOT1L, PLCG1, ITGA6, ITGA5, NTRK1, IGFBP4 |

| | | | | | | |
|---|---|---|---|---|---|---|
| **intracellular signal transduction** | 81 | 76 | 0.9 | 1.4E-04 | CSH1, RAB5B, GRIP1, TUFT1, MAEL, WWC1, PIP5K1B, ARHGAP17, FGF12, CNOT7, PTEN, ARL5A, PAK3, GAB1, NOS3, RAB6B, RAB25, RHOD, DLG5, DEPDC1B, AKT3, NET1, TWIST1, SOCS2, CRYAB, ARTN, TRIM40, ARHGEF12, NEK11, STK3, MAP4K3, HES1, RRAS2, GADD45G, PDE5A, OPHN1, HSPB1, RAB15, SIAH1, TNFAIP1, EPS8L1, GSTP1, KALRN, PRKCZ, CARHSP1, RAB3B, RAP1GAP, RDX, CYTH3, FAM13A, STARD13, RBX1, PSMB5, MTM1, FIS1, AKT1S1, ARHGAP42, TGM2, RHOBTB1, DCX, PHLDA3, RASA1, FBXO8, RHOBTB3, RAB2A, DVL3, CNKSR1, LURAP1, VAV3, EFS, TEAD3, BAD, TAX1BP3, SOD1, TRIM62, CISH, GH2, ICK, RAB36, APBB2, PLEKHA1 | ADCY3, NAF1, ATP1B1, CD8A, IL6ST, EZH2, NCS1, TNFSF15, NFKB1, NFKB2, IL10, SPRY1, MAP3K8, TICAM1, RAPGEF2, IL1A, TNIP3, IRAK2, IRAK1, TNIK, CD3E, NCALD, PDE4D, PATJ, PIK3IP1, TRAT1, DDIT4, NCAM1, CCR7, RND1, SRGAP3, ERN1, SEMA4C, PRNP, EIF2AK3, RASD2, CSF3, CCL2, TNFRSF25, CSF1, DUSP10, KIT, ABCA1, SESN2, TNFRSF4, SRC, LIF, HRH1, IL23A, CCL20, REL, PLCH2, ITGAV, TNFRSF18, ZC3H12A, NFATC2, CD28, PDK1, ITK, FZD8, IL2RB, SPSB1, TP53BP2, IL1RL1, SPHK1, DGKH, ITPR3, ADGRG1, RALGDS, RAB33A, DOT1L, PLCG1, NTRK1, SPTBN1, IGFBP4, SPTAN1 |
| **cell surface receptor signaling pathway** | 67 | 86 | 1.3 | 7.3E-04 | CSH1, SPIN1, PGF, TBX20, AP3S1, PAWR, FGF12, PTEN, DAB2, AES, IFT20, PAK3, GAB1, SPG21, SEMA3B, NOS3, DEPDC1B, RNF146, PID1, SOCS2, STX1B, STK3, HES1, ATP6V1C2, SDC1, DACT2, HSPB1, WASL, ADAMTS3, EMP2, GSTP1, KALRN, PRKCZ, EID2, PPP2R3A, GLIS2, GIPC1, RBX1, PSMB5, P2RY6, AKT1S1, ZNF703, AMER2, SORBS1, TGM2, IL1RAPL2, RASA1, MTMR4, DVL3, CNKSR1, VAV3, ERH, SMAD6, PDK4, BAD, CBY1, TAX1BP3, TRIM62, ANXA4, CISH, IL36RN, GH2, ATP6V1E1, GRK5, ADGRL3, GRB7, PLEKHA1 | NRP2, STIL, CD8A, IL6ST, LGMN, JAG2, TNFSF15, NFKB1, JAG1, SPRY1, CXCR6, TICAM1, UNC5C, RAPGEF2, IL1A, MATK, IRAK2, IRAK1, TNIK, VANGL1, CD3E, TRABD2A, DRAXIN, PDE4D, LDLRAD4, TRAT1, DDIT4, NCAM1, TNFRSF9, ACVR2A, CCR8, CCR7, CCR4, ZFYVE28, SEMA4C, PRNP, EIF2AK3, PMEPA1, KLRC1, CSF3, TRAF1, NKD1, CCL2, TNFRSF25, CXCL3, CSF1, AFAP1L2, KIT, ABCA1, IL7R, TNFRSF4, SRC, LIF, CCL20, ITGB8, ITGAV, CD2, TNFRSF18, ADAMTS10, HSPA5, AXIN2, CD6, NFATC2, CD28, CD7, FZD8, ITK, TCF7, IL2RB, IL1RL1, SPHK1, CTLA4, SMAD3, NLGN2, EVL, IGSF9B, TSPAN18, ADGRG1, ITGA6, PLCG1, ITGA5, NTRK1, SPTBN1, ITGAD, ABL2, IGFBP4 |
| **regulation of intracellular signal transduction** | 48 | 53 | 1.1 | 3.4E-03 | PRKCZ, RAP1GAP, WWC1, PIP5K1B, RDX, ARHGAP17, CYTH3, PTEN, FAM13A, STARD13, FIS1, MTM1, AKT1S1, PAK3, GAB1, ARHGAP42, TGM2, RHOBTB1, RHOD, DEPDC1B, PHLDA3, FBXO8, RASA1, NET1, TWIST1, DVL3, LURAP1, VAV3, SOCS2, TRIM40, BAD, ARHGEF12, SOD1, TRIM62, CISH, STK3, MAP4K3, HES1, PDE5A, GADD45G, OPHN1, HSPB1, SIAH1, GSTP1, EPS8L1, TNFAIP1, PLEKHA1, KALRN | NAF1, CD8A, IL6ST, EZH2, TNFSF15, IL10, SPRY1, MAP3K8, TICAM1, RAPGEF2, IL1A, TNIP3, IRAK2, IRAK1, TNIK, CD3E, PDE4D, PIK3IP1, TRAT1, DDIT4, CCR7, SEMA4C, ERN1, SRGAP3, PRNP, EIF2AK3, RASD2, CSF3, CCL2, TNFRSF25, CSF1, DUSP10, ABCA1, KIT, SESN2, TNFRSF4, SRC, LIF, IL23A, REL, CCL20, TNFRSF18, ZC3H12A, CD28, FZD8, TP53BP2, IL1RL1, SPHK1, ADGRG1, DOT1L, PLCG1, NTRK1, IGFBP4 |
| **(+) regulation of signal transduction** | 37 | 53 | 1.4 | 8.2E-04 | PRKCZ, SPIN1, PPP2R3A, WWC1, GIPC1, STARD10, PTEN, PSMB5, FIS1, DAB2, DYNLL1, SORBS1, PAK3, SMARCB1, GAB1, TGM2, DEPDC1B, RNF146, DVL3, LURAP1, ERH, VAV3, SOCS2, BAD, SOD1, TRIM62, STX1B, STK3, MAP4K3, HES1, ATP6V1C2, PDE5A, GADD45G, SIAH1, ADAMTS3, EMP2, GRB7 | CD8A, IL6ST, EZH2, JAG2, TNFSF15, NFKB1, JAG1, SHE, IL10, SH2D1A, TICAM1, MAP3K8, RAPGEF2, IL1A, IRAK2, IRAK1, TNIK, CD3E, TRAT1, ACVR2A, CCR7, SEMA4C, ERN1, RASD2, CSF3, NKD1, CCL2, TNFRSF25, CSF1, AFAP1L2, KIT, TNFRSF4, SRC, LIF, IL23A, REL, CCL20, TNFRSF18, ZC3H12A, AXIN2, CD28, FZD8, TP53BP2, SPHK1, ASXL1, NLGN2, SMAD3, IGSF9B, ADGRG1, PLCG1, ITGA5, NTRK1, IGFBP4 |
| **(-) regulation of signal transduction** | 36 | 35 | 1.0 | 1.4E-03 | PRKCZ, EID2, PPP2R3A, GLIS2, TBX20, WWC1, PAWR, PTEN, RBX1, PSMB5, MTM1, DAB2, AKT1S1, AES, AMER2, ARHGAP42, NOS3, PHLDA3, RASA1, MTMR4, TWIST1, PID1, DVL3, SOCS2, SMAD6, TRIM40, CBY1, TAX1BP3, STK3, CISH, IL36RN, DACT2, HSPB1, TNFAIP1, GSTP1, PLEKHA1 | NAF1, NKD1, IL6ST, LGMN, EZH2, DUSP10, SESN2, IL10, SRC, LIF, SPRY1, ITGAV, TICAM1, ZC3H12A, HSPA5, AXIN2, IL1A, TNIP3, CD3E, IL1RL1, TRABD2A, DRAXIN, ASXL1, SMAD3, PIK3IP1, RGS16, LDLRAD4, DDIT4, RGS13, ITGA6, ZFYVE28, PPP1R15B, PRNP, PMEPA1, IGFBP4 |

| | | | | | | |
|---|---|---|---|---|---|---|
| (+) regulation of cell communication | 44 | 54 | 1.2 | 4.7E-04 | SYT1, PRKCZ, RAB3B, SPIN1, PPP2R3A, ANO1, WWC1, GIPC1, STARD10, PTEN, PSMB5, FIS1, DAB2, DYNLL1, SORBS1, ANK3, SMARCB1, PAK3, GAB1, TGM2, DEPDC1B, RNF146, DVL3, LURAP1, ERH, VAV3, STX3, SOCS2, BAD, SOD1, TRIM62, STX1B, STK3, MAP4K3, HES1, ATP6V1C2, VAMP8, PDE5A, GADD45G, SIAH1, ADAMTS3, EMP2, GRB7, KALRN | CD8A, IL6ST, EZH2, JAG2, TNFSF15, NFKB1, JAG1, SHE, IL10, SH2D1A, MAP3K8, TICAM1, RAPGEF2, IL1A, IRAK2, IRAK1, TNIK, CD3E, TRAT1, ACVR2A, CCR7, SEMA4C, ERN1, RASD2, CSF3, NKD1, CCL2, TNFRSF25, CSF1, AFAP1L2, KIT, TNFRSF4, SRC, LIF, IL23A, REL, CCL20, TNFRSF18, ZC3H12A, AXIN2, CD28, FZD8, TP53BP2, SPHK1, ASXL1, NLGN2, SMAD3, IGSF9B, ITPR3, ADGRG1, PLCG1, ITGA5, NTRK1, IGFBP4 |
| (-) regulation of cell communication | 36 | 36 | 1.0 | 6.7E-03 | PRKCZ, EID2, PPP2R3A, GLIS2, TBX20, WWC1, PAWR, PTEN, RBX1, PSMB5, MTM1, DAB2, AKT1S1, AES, AMER2, ARHGAP42, NOS3, PHLDA3, RASA1, MTMR4, TWIST1, PID1, DVL3, SOCS2, SMAD6, TRIM40, CBY1, TAX1BP3, STK3, CISH, IL36RN, DACT2, HSPB1, TNFAIP1, GSTP1, PLEKHA1 | NAF1, NKD1, IL6ST, LGMN, EZH2, DUSP10, SESN2, IL10, SRC, LIF, SPRY1, ITGAV, TICAM1, ZC3H12A, HSPA5, AXIN2, IL1A, TNIP3, CD3E, IL1RL1, TRABD2A, DRAXIN, ASXL1, SMAD3, ASIC1, PIK3IP1, RGS16, LDLRAD4, DDIT4, RGS13, ITGA6, ZFYVE28, PPP1R15B, PRNP, PMEPA1, IGFBP4 |
| phosphate-containing compound metabolic process | 91 | 86 | 0.9 | 2.2E-04 | MOCOS, ALPPL2, IMPA2, GDA, WWC1, PIP5K1B, PPCS, SULT2B1, FGF12, PTEN, PIFO, DAB2, DYNLL1, PAK3, GAB1, NOS3, PPP1R14C, PPP1R14B, AKT3, TWIST1, PID1, CEP85, SOCS2, CRYAB, PKIG, TTC7B, ARTN, LGALS13, COQ9, CAMSAP3, PKIB, NEK11, STK3, NME6, MAP4K3, HES1, UMPS, RFK, ADK, PUDP, GADD45G, PGM1, PDE5A, HSPB1, MAPRE3, EMP2, GSTP1, NEK7, KALRN, SSU72, PRKCZ, PPP2R3A, NDUFB7, GNE, SSH3, ABHD5, MMD, ALPP, SMUG1, RBX1, PSMB5, MTM1, PFN2, AKT1S1, REXO2, MLLT1, UCK2, DCX, MTMR7, RASA1, MTMR4, DVL3, PLA2G16, VAV3, ERH, CAP2, NDUFA6, SMAD6, PDK4, PPP1R11, AK3, BPGM, BAD, SOD1, CISH, ICK, CHCHD10, AKTIP, GFPT2, BTBD10, GRK5 | ADCY3, NAF1, ATP1B1, CDK17, IL6ST, EZH2, TNFSF15, NFKB1, SPRY1, MAP3K8, RANBP2, AGPAT4, RAPGEF2, CDK15, RAMP1, IL1A, MATK, IRAK2, IRAK1, TNIK, CAMK1G, CD3E, PGAP1, LDB2, PDE4D, PIK3IP1, LDLRAD4, TRAT1, DDIT4, NCAM1, ACVR2A, CCR7, CCND2, ZFYVE28, ERN1, SEMA4C, PRNP, PPP1R15B, EIF2AK3, CLN8, PMEPA1, CSF3, CCL2, TNFRSF25, ENPP3, CSF1, DUSP10, HK2, AFAP1L2, KIT, ABCA1, SESN2, TNFRSF4, SRC, LIF, HRH1, TSPYL2, IL23A, LPCAT1, CCL20, PLCH2, ITGAV, ENO2, TNFRSF18, ZC3H12A, HSPA5, AXIN2, EHD4, CD28, PDK1, FZD8, ITK, IL2RB, MEX3B, SPHK1, SMAD3, DGKH, PLCG1, ITGA6, ITGA5, NTRK1, SPTBN1, ADM2, ABL2, IGFBP4, SPTAN1 |
| regulation of phosphorus metabolic process | 42 | 59 | 1.4 | 4.4E-04 | PRKCZ, WWC1, MMD, PTEN, PIFO, DAB2, PFN2, AKT1S1, DYNLL1, PAK3, GAB1, MLLT1, NOS3, PPP1R14C, PPP1R14B, TWIST1, PID1, CEP85, DVL3, CAP2, VAV3, SOCS2, SMAD6, PKIG, PPP1R11, PKIB, BPGM, CAMSAP3, BAD, SOD1, STK3, CISH, MAP4K3, HES1, AKTIP, PDE5A, GADD45G, BTBD10, HSPB1, EMP2, MAPRE3, GSTP1 | ADCY3, NAF1, IL6ST, EZH2, TNFSF15, SPRY1, MAP3K8, RANBP2, RAPGEF2, RAMP1, IL1A, IRAK2, IRAK1, TNIK, CD3E, LDB2, PDE4D, PIK3IP1, LDLRAD4, DDIT4, ACVR2A, CCR7, CCND2, ZFYVE28, SEMA4C, ERN1, PRNP, PPP1R15B, EIF2AK3, PMEPA1, CSF3, CCL2, TNFRSF25, CSF1, DUSP10, AFAP1L2, ABCA1, KIT, SESN2, TNFRSF4, SRC, LIF, HRH1, IL23A, TSPYL2, CCL20, TNFRSF18, ZC3H12A, HSPA5, AXIN2, EHD4, FZD8, SPHK1, SMAD3, PLCG1, ITGA6, ITGA5, NTRK1, IGFBP4 |
| (+) regulation of phosphorus metabolic process | 25 | 43 | 1.7 | 2.0E-03 | PRKCZ, MMD, WWC1, PTEN, PIFO, PFN2, DAB2, PAK3, GAB1, NOS3, PID1, DVL3, VAV3, CAP2, BAD, SOD1, STK3, HES1, MAP4K3, AKTIP, GADD45G, PDE5A, BTBD10, MAPRE3, EMP2 | CSF3, ADCY3, CCL2, IL6ST, TNFRSF25, CSF1, EZH2, TNFSF15, AFAP1L2, ABCA1, KIT, TNFRSF4, SRC, LIF, HRH1, IL23A, CCL20, MAP3K8, TNFRSF18, ZC3H12A, HSPA5, AXIN2, RAPGEF2, RAMP1, IL1A, EHD4, IRAK2, IRAK1, FZD8, TNIK, CD3E, SPHK1, SMAD3, ACVR2A, CCR7, ITGA6, PLCG1, CCND2, ITGA5, NTRK1, SEMA4C, ERN1, IGFBP4 |

| | | | | | | |
|---|---|---|---|---|---|---|
| **cell differentiation** | 106 | 86 | 0.8 | 7.9E-03 | SYT1, GLDN, PGF, MAEL, POSTN, ILDR2, TPD52, DAB2, KDF1, IFT20, ANK3, CCSAP, BTBD3, RAB25, EIF2B2, FBXO22, NET1, TWIST1, PID1, CECR2, CHODL, STK3, NEBL, HES1, CRCT1, PDE5A, OPHN1, EMP2, KALRN, KAZN, EID2, PPP2R3A, SSH3, MMD, PAQR7, UBE2V2, ECE2, SMARCB1, NGRN, TMEM79, MAFF, PARD6B, SMAD6, MEA1, BPGM, TEAD3, TRIM62, THSD7A, HOPX, GRK5, PLEKHA1, STEAP4, HLF, HTATIP2, NIF3L1, GRIP1, POU6F2, TBX20, PTEN, BZW2, EFHD1, PAK3, CAMSAP2, SEMA3B, DLG5, ZFAT, KIF2A, TCHH, STX3, KIF17, MPP5, ARTN, NECTIN3, CAMSAP3, STX1B, SDC1, DACT2, CLIC5, RRAS2, GADD45G, SCIN, SIAH1, GSTP1, STON2, PRKCZ, RAP1GAP, FHL1, GLIS2, ABHD5, RDX, CBR1, ZNF703, ZNF750, DCX, ETV5, RASA1, ETV4, ERH, TBX3, EXPH5, BAD, CBY1, SOD1, ANXA4, APBB2, ADGRL3 | NRP2, FOSL2, CD8A, IL6ST, EZH2, JAG2, NCS1, SLFN5, NFKB1, JAG1, NFKB2, SLC7A5, IL10, NOCT, S1PR1, PBXIP1, SNPH, SLC9B2, UNC5C, RAPGEF2, IL1A, MATK, TNIK, CD3E, DRAXIN, MMP19, ZHX2, NECTIN2, PDE4D, LDLRAD4, DDIT4, NCAM1, ACVR2A, CCR7, RND1, CCR4, SEMA4C, EIF2AK3, KDM6B, CSF3, PHLDB1, NKD1, CCL2, CSF1, DUSP10, MYEF2, KIT, ABCA1, IL7R, SRC, MLF1, ITM2A, LIF, LAMB3, IL23A, ITGAV, BCL11B, CD2, ZC3H12A, HEG1, COL6A1, NFATC2, AXIN2, ETV3, CD28, MAF, FZD8, ITK, TCF7, ASXL1, TMEM120B, CTLA4, CENPF, SMAD3, EVL, ADGRG1, WHRN, ABCG1, PENK, ITGA6, ITGA5, NTRK1, SPTBN1, ABL2, SPTAN1, FEZ1 |
| **system development** | 127 | 98 | 0.8 | 3.9E-03 | SYT1, CGA, GDA, GLDN, PGF, TUFT1, POSTN, ILDR2, FGF12, MYLIP, TPD52, DAB2, KDF1, IFT20, ANK3, CCSAP, BTBD3, GAB1, TRIM45, PMS2, EIF2B2, FBXO22, TWIST1, CRYAB, CECR2, CHODL, CDO1, STK3, NEBL, HES1, CRCT1, TFAP2A, HSPB1, OPHN1, EMP2, KALRN, KAZN, EID2, PPP2R3A, SSH3, MMD, UBE2V2, PSMB5, ECE2, SMARCB1, NGRN, IL1RAPL2, TMEM79, CYP19A1, DVL3, MAFF, PARD6B, VAV3, CRIP2, SMAD6, MEA1, BPGM, TEAD3, TRIM62, LIN7A, THSD7A, HOPX, PLEKHA1, BMI1, HLF, HTATIP2, NIF3L1, GRIP1, POU6F2, TBX20, PTEN, BZW2, EFHD1, AES, DYNLL1, PAK3, PCP4, CAMSAP2, NOS3, SEMA3B, STRA6, DLG5, ZFAT, COX17, RS1, KIF2A, TCHH, STX3, MAN1A2, KIF17, MPP5, ARTN, CAMSAP3, NECTIN3, STX1B, SDC1, UMPS, DACT2, CLIC5, SCIN, SIAH1, GSTP1, STON2, PRKCZ, GLRX5, RAP1GAP, HSD17B1, FHL1, GLIS2, STARD13, MTM1, NPAS2, ZNF703, TGM2, PAFAH1B2, DCX, ETV5, ETV4, RASA1, TBX3, PLAC1, EXPH5, CBY1, BAD, SOD1, APBB2, ADGRL3 | NRP2, STIL, FOSL2, CD8A, IL6ST, EZH2, JAG2, NCS1, JAG1, NFKB2, SLC7A5, IL10, SPRY1, S1PR1, SNPH, SLC9B2, UNC5C, RAPGEF2, RAMP1, IL1A, TNIK, VANGL1, CD3E, PGAP1, DRAXIN, MMP19, ZHX2, LDB2, LDLRAD4, DDIT4, NCAM1, AMIGO2, ACVR2A, CCR7, RND1, XIRP1, CCR4, SEMA4C, EIF2AK3, CLN8, KDM6B, CSF3, ABLIM1, NKD1, CCL2, SPOCK2, CSF1, HK2, DUSP10, NINJ1, MYEF2, KIT, IL7R, SRC, MLF1, ITM2A, LIF, IL23A, LPCAT1, ITGB8, ICOS, ITGAV, BCL11B, CD2, ZC3H12A, HEG1, HSPA5, AXIN2, BCOR, CD28, MAF, FZD8, ITK, TCF7, ASXL1, SPHK1, CTLA4, CENPF, SMAD3, NLGN2, EVL, IGSF9B, ADGRG1, WHRN, PENK, ITGA6, PLCG1, ITGA5, BNC2, NTRK1, MAMLD1, SPTBN1, ADM2, TMEM41B, ABL2, IGFBP4, SPTAN1, FEZ1 |
| **nervous system development** | 67 | 55 | 0.8 | 5.7E-03 | SYT1, GDA, GLDN, NIF3L1, GRIP1, POU6F2, TBX20, POSTN, MYLIP, FGF12, PTEN, BZW2, EFHD1, DYNLL1, IFT20, PCP4, PAK3, ANK3, CAMSAP2, CCSAP, BTBD3, SEMA3B, DLG5, EIF2B2, COX17, KIF2A, TWIST1, STX3, KIF17, MPP5, ARTN, CHODL, CECR2, CAMSAP3, STX1B, STK3, HES1, CLIC5, TFAP2A, OPHN1, SIAH1, GSTP1, KALRN, PRKCZ, PPP2R3A, RAP1GAP, SSH3, GLIS2, MMD, UBE2V2, NPAS2, ECE2, SMARCB1, NGRN, PAFAH1B2, DCX, IL1RAPL2, ETV5, ETV4, PARD6B, DVL3, TBX3, TEAD3, BAD, SOD1, APBB2, ADGRL3 | NRP2, STIL, IL6ST, EZH2, JAG2, NCS1, JAG1, SLC7A5, S1PR1, SNPH, UNC5C, RAPGEF2, TNIK, DRAXIN, PGAP1, ZHX2, DDIT4, NCAM1, AMIGO2, RND1, CCR4, SEMA4C, CLN8, EIF2AK3, KDM6B, NKD1, CCL2, SPOCK2, CSF1, DUSP10, MYEF2, NINJ1, KIT, SRC, ITM2A, LIF, BCL11B, ZC3H12A, HSPA5, FZD8, TCF7, SPHK1, NLGN2, CENPF, EVL, IGSF9B, WHRN, ADGRG1, PENK, NTRK1, SPTBN1, TMEM41B, ABL2, FEZ1, SPTAN1 |
| **(+) regulation of synaptic transmission** | 7 | 5 | 0.7 | 9.6E-03 | PRKCZ, SYT1, RAB3B, STX3, STX1B, PTEN, KALRN | CCL2, NTRK1, NLGN2, IGSF9B, ITPR3 |

| | | | | | | |
|---|---|---|---|---|---|---|
| **single-organism organelle organization** | 68 | 28 | 0.4 | 3.1E-03 | SYT1, CHMP5, PEX3, MYLIP, PAWR, TMEM141, AUNIP, ANKRD53, DYNLL1, IFT20, PAK3, ANK3, CAMSAP2, CCSAP, RHOD, KIF2A, PID1, CEP85, STX3, DSN1, CRYAB, VTI1B, CAMSAP3, SPIRE2, STX1B, NEBL, VAMP8, SCIN, OPHN1, WASL, EMP2, TNFAIP1, MAP7D3, NEK7, TPPP3, PRKCZ, STX7, CRIPT, SSH3, BET1, RDX, CHMP2B, FIS1, MTM1, PFN2, PEX19, SORBS1, FIGN, C10ORF90, SKA2, SNAP23, RASA1, SYNPO, CAPN6, GABARAPL1, LURAP1, VAV3, CAP2, MSRB1, BAD, CBY1, SOD1, MSRB2, ICK, SLAIN2, HEBP2, CCDC113, KCTD17 | ABLIM1, CSF3, STIL, PRC1, HK2, KIT, TPM2, SRC, SPRY1, SLC16A1, S1PR1, STARD9, TBC1D4, RANBP2, AXIN2, TNIK, TP53BP2, SMAD3, CENPF, NECTIN2, EVL, CCR7, RND1, XIRP1, SPAG5, SPTBN1, ABL2, SPTAN1 |
| **cytoskeleton organization** | 47 | 25 | 0.5 | 3.0E-03 | TPPP3, PRKCZ, CHMP5, CRIPT, SSH3, RDX, PAWR, MYLIP, TMEM141, AUNIP, CHMP2B, MTM1, PFN2, ANKRD53, DYNLL1, SORBS1, FIGN, ANK3, PAK3, CCSAP, CAMSAP2, SKA2, RHOD, RASA1, KIF2A, SYNPO, CAPN6, CEP85, LURAP1, CAP2, CRYAB, MSRB1, CECR2, CAMSAP3, SPIRE2, SOD1, MSRB2, NEBL, SLAIN2, SCIN, TUBAL3, OPHN1, WASL, EMP2, MAP7D3, TNFAIP1, NEK7 | ABLIM1, CSF3, PHLDB1, STIL, CCL2, PRC1, KIT, TPM2, SRC, SPRY1, SLC16A1, S1PR1, STARD9, TUBB6, TNIK, SMAD3, NECTIN2, EVL, CCR7, RND1, XIRP1, SPAG5, SPTBN1, ABL2, SPTAN1 |
| **plasma membrane organization** | 20 | 6 | 0.3 | 7.1E-04 | PID1, PRKCZ, STX3, STX7, TTC7B, MPP5, VTI1B, LYPLA1, RDX, SOD1, PTEN, DAB2, SORBS1, PACSIN3, VAMP8, ANK3, IFT20, WASL, EMP2, KALRN | ATP1B1, TNIK, GOLGA7B, SPTBN1, RAPGEF2, RAMP1 |
| **regulation of plasma membrane organization** | 10 | 1 | 0.1 | 1.8E-03 | PID1, DAB2, STX7, STX3, SORBS1, VAMP8, VTI1B, LYPLA1, WASL, KALRN | SPTBN1 |

**Supplemental Table 5. Differential expression of M1/M2 markers in MIMs vs. HBCs.** A panel of 19 anti-inflammatory (M1) and 21 pro-inflammatory (M2) markers were identified based on a broad literature review. Out of this panel, 24 genes were observed to be differentially expressed between MIMs and HBCs (average expression>0, p<0.01 and absolute $log_2FC > 1$; please note less conservative criteria than unbiased analysis). Abbreviations include: $Log_2$ average counts in MIMs (MIM_Ct) or HBCs (HBC_Ct); fold change difference between MIMs and HBCs ($log_2FC$); and significance (p-value). Red color indicates significance.

| Gene | Function | Status | MIM_Ct | HBC_Ct | $log_2FC$ | p-value |
|---|---|---|---|---|---|---|
| **CCL2** | **Chemokine** | **M1** | **1.7** | **5.3** | **-3.6** | **6.3E-06** |
| CCL8 | Chemokine | M1 | -0.5 | 2.6 | -3.2 | 3.5E-04 |
| **CCR7** | **Chemokine Receptor** | **M1** | **2.9** | **7.7** | **-4.8** | **5.1E-07** |
| **CD80** | **Membrane Receptor** | **M1** | **0.4** | **3.7** | **-3.3** | **7.4E-05** |
| **CXCL1** | **Chemokine** | **M1** | **4.3** | **6.6** | **-2.3** | **6.1E-04** |
| **CXCL2** | **Chemokine** | **M1** | **5.5** | **9.5** | **-3.9** | **1.0E-04** |
| **CXCL3** | **Chemokine** | **M1** | **4.2** | **8.9** | **-4.6** | **5.5E-06** |
| **CXCL5** | **Chemokine** | **M1** | **1.5** | **3.5** | **-2.0** | **3.6E-04** |
| **CXCL8** | **Chemokine** | **M1** | **9.0** | **13.0** | **-4.0** | **9.8E-05** |
| CXCL9 | Chemokine | M1 | 2.8 | 4.7 | -1.8 | 1.3E-01 |
| CXCL10 | Chemokine | M1 | 2.4 | 4.3 | -1.7 | 1.1E-01 |
| **IL1A** | **Cytokine** | **M1** | **2.3** | **8.4** | **-5.9** | **4.4E-06** |
| **IL1B** | **Cytokine** | **M1** | **9.1** | **12.4** | **-3.4** | **1.6E-03** |
| **IL6** | **Cytokine** | **M1** | **4.2** | **7.8** | **-3.5** | **2.4E-03** |
| IL12A | Cytokine | M1 | -1.8 | 0.0 | -1.7 | 1.5E-02 |
| IL12B | Cytokine | M1 | -0.4 | -1.3 | 0.9 | 1.5E-01 |
| **IL23A** | **Cytokine** | **M1** | **0.3** | **6.5** | **-6.2** | **2.3E-06** |
| TLR2 | Toll-Like Receptor | M1 | 6.0 | 8.1 | -2.1 | 1.1E-02 |
| TLR4 | Toll-Like Receptor | M1 | 5.6 | 7.7 | -2.1 | 1.1E-02 |
| **CCL22** | **Chemokine** | **M2** | **0.2** | **4.3** | **-4.0** | **4.9E-04** |
| CCL24 | Chemokine | M2 | -2.3 | -1.9 | -0.3 | 7.3E-01 |
| CCR2 | Chemokine Receptor | M2 | 4.1 | 2.4 | 1.7 | 1.0E-01 |
| **CD163** | **Scavenger Receptor** | **M2** | **6.7** | **8.9** | **-2.2** | **1.3E-03** |
| **CD209** | **Membrane Receptor** | **M2** | **1.6** | **4.3** | **-2.8** | **2.3E-04** |
| CXCR1 | Chemokine Receptor | M2 | 1.6 | 2.9 | -1.4 | 3.4E-01 |
| CXCR2 | Chemokine Receptor | M2 | 2.6 | 3.8 | -1.3 | 3.3E-01 |
| **FOLR2** | **Folate Receptor** | **M2** | **1.9** | **5.3** | **-3.3** | **5.6E-04** |
| **IL1R2** | **Cytokine Receptor** | **M2** | **3.1** | **5.4** | **-2.3** | **2.3E-03** |
| **IL4R** | **Cytokine Receptor** | **M2** | **5.8** | **8.2** | **-2.3** | **1.0E-03** |
| **IL10** | **Cytokine** | **M2** | **2.1** | **5.8** | **-3.7** | **7.1E-06** |
| PDGFA | Growth Factor | M2 | 1.7 | 2.5 | -0.9 | 2.9E-02 |
| **PDGFB** | **Growth Factor** | **M2** | **5.2** | **6.4** | **-1.2** | **2.1E-03** |
| **PDGFC** | **Growth Factor** | **M2** | **1.0** | **3.7** | **-2.6** | **4.5E-04** |
| **PDGFD** | **Growth Factor** | **M2** | **1.1** | **3.2** | **-2.1** | **1.0E-04** |
| TGFB1 | Growth Factor | M2 | 8.9 | 9.2 | -0.3 | 3.0E-01 |
| TGFB2 | Growth Factor | M2 | -0.1 | 3.3 | -3.3 | 3.1E-03 |
| TGFB3 | Growth Factor | M2 | -1.5 | 0.8 | -2.3 | 1.3E-03 |
| **VEGFA** | **Growth Factor** | **M2** | **5.5** | **9.7** | **-4.2** | **1.0E-05** |
| **VEGFB** | **Growth Factor** | **M2** | **6.1** | **5.0** | **1.0** | **2.0E-03** |
| VEGFC | Growth Factor | M2 | -3.2 | -1.1 | -2.1 | 8.5E-02 |

**Supplemental Table 6: Gene ontology analysis of genes Influenced by gravidity in MIMs or HBCs.** Significant biological processes influenced by gravid status in MIMs or HBCs identified via DAVID. Criteria: p<0.01 in either MIMs or HBCs and minimum differentially expressed genes ≥ 10.

| Term | MIMs (p) | MIMs (# genes) | HBCs (p) | HBCs (# genes) | MIMs (genes) | HBCs (genes) |
|---|---|---|---|---|---|---|
| **system development** | 0.014 | 47 | 0.003 | 45 | E2F1, PALM, TNF, TCAP, POMK, E2F7, EZH2, DUOX2, CABP4, CKB, FAM83D, PACSIN1, KRT27, FRMD7, BOK, NRARP, RASGRP1, TNR, IL1B, TNFRSF19, SIK1, THBS1, FOSL1, KNDC1, IL1A, HELLS, ICAM1, AR, IL6, MKI67, CHAC1, IL1RL2, MACROD2, NR4A3, UPK3A, NLRP3, MSC, DHRS2, GNGT1, MYO18B, KRT17, WDR62, IFNB1, HES4, F3, SPTBN2, MYLK | CSF3, CSF2, STOX1, CPLX2, SLC38A3, WNT5B, HNF1A, RBP1, IGF2BP1, LRRC17, DDR2, ADCYAP1, VCAM1, SLC1A2, HOXA3, CYP27B1, HEY1, CDNF, ROBO1, IFNG, MT1G, PLAG1, UCN, PTPRD, GNAO1, RBM20, MACROD2, RPGRIP1, UPK3A, COL5A3, HMGA2, SIGLEC15, MSC, TNNI1, SSTR2, RND1, BVES, NUPR1, EPGN, MEOX1, TENM2, LRP6, MYRF, PLA2G2D, ADAMTS4 |
| **animal organ development** | 0.012 | 37 | 0.003 | 36 | E2F1, TNF, TCAP, POMK, E2F7, EZH2, DUOX2, CABP4, CKB, FAM83D, KRT27, BOK, NRARP, RASGRP1, TNR, TNFRSF19, SIK1, FOSL1, KNDC1, ICAM1, AR, IL6, MKI67, IL1RL2, MACROD2, NR4A3, UPK3A, NLRP3, MSC, GNGT1, DHRS2, MYO18B, KRT17, WDR62, IFNB1, SPTBN2, MYLK | CSF3, CSF2, STOX1, SLC38A3, WNT5B, HNF1A, RBP1, IGF2BP1, LRRC17, DDR2, ADCYAP1, VCAM1, SLC1A2, HOXA3, CYP27B1, HEY1, ROBO1, IFNG, MT1G, PLAG1, GNAO1, RBM20, MACROD2, RPGRIP1, UPK3A, HMGA2, COL5A3, MSC, SIGLEC15, TNNI1, SSTR2, BVES, NUPR1, MEOX1, LRP6, PLA2G2D |
| **regulation of signal transduction** | 0.001 | 38 | 0.098 | 26 | E2F1, TRAF1, PALM, TNF, EZH2, SPINK1, FAM83D, KIF7, JSRP1, TRIM68, CCL20, FRMD7, BOK, NRARP, RASGRP1, APOC3, TNFRSF19, IL1B, GNG4, THBS1, HELLS, IL1A, ICAM1, AR, IL6, IL1RL1, CHAC1, TPX2, RPH3AL, BIRC5, NLRP3, GNGT1, LYNX1, IFNB1, F3, DUSP8, GADD45A, PTGDR2 | CSF3, CSF2, STOX1, FGFR4, HNF1A, WNT5B, NKD2, ARHGEF26, TNFSF15, BDKRB2, ADCYAP1, AKR1C2, CYP27B1, HEY1, JSRP1, ROBO1, IFNG, ARC, DGKI, HMGA2, ARHGAP32, CCL13, EPGN, LRP6, RWDD3, RASD2 |
| **intracellular signal transduction** | 0.005 | 34 | 0.172 | 24 | E2F1, TNF, E2F7, EZH2, SPINK1, FAM83D, CCL20, FRMD7, JSRP1, GRIN2C, BOK, RASGRP1, APOC3, TNFRSF19, IL1B, SIK1, THBS1, KNDC1, HELLS, IL1A, ICAM1, AR, IL6, CHAC1, IL1RL1, TPX2, CDC25C, NLRP3, IFNB1, F3, SPTBN2, PTGDR2, DUSP8, GADD45A | CSF3, CSF2, STOX1, FGFR4, ARHGEF26, TNFSF15, BDKRB2, DGKI, HMGA2, ADCYAP1, VCAM1, TIFAB, AKR1C2, ARHGAP32, CCL13, RND1, NUPR1, JSRP1, EPGN, ROBO1, TENM2, IFNG, SELE, RASD2 |
| **regulation of intracellular signal transduction** | 0.004 | 25 | 0.095 | 18 | TNF, EZH2, SPINK1, FAM83D, FRMD7, CCL20, JSRP1, BOK, RASGRP1, APOC3, TNFRSF19, IL1B, THBS1, HELLS, IL1A, ICAM1, IL6, AR, IL1RL1, TPX2, NLRP3, IFNB1, F3, DUSP8, GADD45A | CSF3, CSF2, STOX1, FGFR4, ARHGEF26, TNFSF15, DGKI, BDKRB2, HMGA2, ADCYAP1, AKR1C2, ARHGAP32, CCL13, JSRP1, EPGN, ROBO1, IFNG, RASD2 |
| **negative regulation of cell communication** | 0.006 | 19 | 0.459 | 10 | ICAM1, PALM, AR, IL6, TNF, CHAC1, IL1RL1, EZH2, RPH3AL, SPINK1, NLRP3, KIF7, NRARP, TNR, IL1B, THBS1, DUSP8, HELLS, IL1A | CSF2, UCN, WNT5B, NKD2, HEY1, ROBO1, LRP6, ASIC1, BDKRB2, HMGA2 |

| | | | | | | |
|---|---|---|---|---|---|---|
| **negative regulation of signal transduction** | 0.006 | 18 | 0.657 | 8 | ICAM1, PALM, AR, IL6, TNF, CHAC1, IL1RL1, EZH2, RPH3AL, SPINK1, NLRP3, KIF7, NRARP, IL1B, THBS1, DUSP8, IL1A, HELLS | CSF2, WNT5B, NKD2, HEY1, ROBO1, LRP6, BDKRB2, HMGA2 |
| **signal transduction by protein phosphorylation** | 0.004 | 16 | 0.873 | 5 | ICAM1, AR, IL6, TNF, EZH2, FAM83D, CCL20, GRIN2C, RASGRP1, SPTBN2, TNFRSF19, IL1B, THBS1, GADD45A, DUSP8, IL1A | CSF2, CCL13, FGFR4, EPGN, ADCYAP1 |
| **cell-cell signaling** | 0.675 | 12 | 0.010 | 20 | AR, IL6, TNF, CCL20, NRARP, GRIN2C, TNR, RPH3AL, IL1B, BIRC5, PTGDR2, KCNJ3 | ARC, PTPRD, UCN, CPLX2, NKD2, WNT5B, HNF1A, GNAO1, SLC6A12, DGKI, ASIC1, HMGA2, ADCYAP1, SSTR2, CCL13, SLC1A2, KCNQ3, IFNG, LRP6, RASD2 |
| **regulation of protein secretion** | 0.001 | 11 | 0.940 | 2 | IL6, TNF, IL1RL1, RASGRP1, RPH3AL, IL1B, BIRC5, NLRP3, NLRP10, IL1A, SCAMP5 | HNF1A, IFNG |
| **protein secretion** | 0.004 | 11 | 0.966 | 2 | IL6, TNF, IL1RL1, RASGRP1, RPH3AL, IL1B, BIRC5, NLRP3, NLRP10, IL1A, SCAMP5 | HNF1A, IFNG |
| **positive regulation of secretion** | 0.002 | 10 | 0.460 | 4 | IL6, TNF, IL1RL1, RASGRP1, RPH3AL, IL1B, NLRP3, NLRP10, IL1A, SCAMP5 | UCN, P2RY2, IFNG, ADCYAP1 |
| **positive regulation of secretion by cell** | 0.001 | 10 | 0.676 | 3 | IL6, TNF, IL1RL1, RASGRP1, RPH3AL, IL1B, NLRP3, NLRP10, IL1A, SCAMP5 | UCN, IFNG, ADCYAP1 |
| **positive regulation of protein secretion** | 0.000 | 10 | 1.000 | 1 | IL6, TNF, IL1RL1, RASGRP1, RPH3AL, IL1B, NLRP3, NLRP10, IL1A, SCAMP5 | IFNG |
| **synaptic signaling** | 0.865 | 4 | 0.000 | 14 | GRIN2C, TNR, RPH3AL, PTGDR2 | ARC, UCN, CPLX2, PTPRD, SLC6A12, ASIC1, DGKI, ADCYAP1, SLC1A2, SSTR2, KCNQ3, IFNG, LRP6, RASD2 |
| **regulation of protein metabolic process** | 0.008 | 32 | 0.412 | 20 | TNF, EZH2, SPINK1, PTTG1, FAM83D, CCL20, GRIN2C, BOK, RASGRP1, TNFRSF19, IL1B, THBS1, KNDC1, IL1A, RNF144A, ICAM1, AR, IL6, ASTL, CHAC1, PTTG3P, TPX2, BIRC5, CDC25C, NLRP3, KRT17, IFNB1, F3, BUB1B, WFDC1, DUSP8, GADD45A | CSF3, CSF2, STOX1, FGFR4, UCN, NKD2, TNFSF15, IGF2BP1, BDKRB2, AZIN2, DDR2, ADCYAP1, CCL13, NUPR1, EPGN, ROBO1, IFNG, LRP6, RWDD3, RASD2 |
| **regulation of cellular protein metabolic process** | 0.005 | 31 | 0.392 | 19 | TNF, EZH2, SPINK1, PTTG1, FAM83D, CCL20, BOK, RASGRP1, TNFRSF19, IL1B, THBS1, KNDC1, IL1A, RNF144A, ICAM1, AR, IL6, ASTL, CHAC1, PTTG3P, TPX2, BIRC5, CDC25C, NLRP3, KRT17, IFNB1, F3, BUB1B, WFDC1, DUSP8, GADD45A | CSF3, CSF2, STOX1, FGFR4, UCN, NKD2, TNFSF15, IGF2BP1, BDKRB2, DDR2, ADCYAP1, CCL13, NUPR1, EPGN, ROBO1, IFNG, LRP6, RWDD3, RASD2 |
| **positive regulation of cellular protein metabolic process** | 0.004 | 22 | 0.075 | 16 | RNF144A, ICAM1, AR, IL6, TNF, ASTL, EZH2, TPX2, NLRP3, CCL20, KRT17, BOK, IFNB1, F3, RASGRP1, TNFRSF19, IL1B, BUB1B, THBS1, GADD45A, KNDC1, IL1A | CSF3, CSF2, STOX1, FGFR4, UCN, NKD2, TNFSF15, DDR2, ADCYAP1, CCL13, NUPR1, EPGN, ROBO1, IFNG, RWDD3, RASD2 |
| **positive regulation of protein metabolic process** | 0.008 | 22 | 0.111 | 16 | RNF144A, ICAM1, AR, IL6, TNF, ASTL, EZH2, TPX2, NLRP3, CCL20, KRT17, BOK, IFNB1, F3, RASGRP1, TNFRSF19, IL1B, BUB1B, THBS1, GADD45A, KNDC1, IL1A | CSF3, CSF2, STOX1, FGFR4, UCN, NKD2, TNFSF15, DDR2, ADCYAP1, CCL13, NUPR1, EPGN, ROBO1, IFNG, RWDD3, RASD2 |

| | | | | | | |
|---|---|---|---|---|---|---|
| **MAPK cascade** | 0.003 | 16 | 0.852 | 5 | ICAM1, AR, IL6, TNF, EZH2, FAM83D, CCL20, GRIN2C, RASGRP1, SPTBN2, TNFRSF19, IL1B, THBS1, GADD45A, DUSP8, IL1A | CSF2, CCL13, FGFR4, EPGN, ADCYAP1 |
| **programmed cell death** | 0.003 | 27 | 0.856 | 11 | TRAF1, LGALS17A, E2F1, TNF, BOK, TNFRSF19, IL1B, THBS1, SIK1, FOSL1, HELLS, IL1A, ICAM1, AR, IL6, CHAC1, TPX2, BIRC5, NR4A3, MCM2, NLRP3, GNGT1, DHRS2, IFNB1, F3, BUB1B, GADD45A | KCNMA1, CSF2, UCN, NUPR1, ROBO1, IFNG, TNFSF15, LRP6, BDKRB2, HMGA2, ADCYAP1 |
| **regulation of programmed cell death** | 0.008 | 21 | 0.546 | 11 | E2F1, TRAF1, LGALS17A, ICAM1, AR, IL6, TNF, BIRC5, NR4A3, NLRP3, DHRS2, IFNB1, BOK, F3, IL1B, THBS1, SIK1, GADD45A, FOSL1, HELLS, IL1A | KCNMA1, CSF2, UCN, NUPR1, ROBO1, IFNG, TNFSF15, LRP6, BDKRB2, HMGA2, ADCYAP1 |
| **positive regulation of programmed cell death** | 0.001 | 14 | 0.387 | 6 | LGALS17A, E2F1, ICAM1, IL6, TNF, NR4A3, NLRP3, IFNB1, BOK, F3, THBS1, SIK1, GADD45A, FOSL1 | KCNMA1, NUPR1, ROBO1, IFNG, TNFSF15, HMGA2 |
| **positive regulation of cell death** | 0.001 | 14 | 0.429 | 6 | LGALS17A, E2F1, ICAM1, IL6, TNF, NR4A3, NLRP3, IFNB1, BOK, F3, THBS1, SIK1, GADD45A, FOSL1 | KCNMA1, NUPR1, ROBO1, IFNG, TNFSF15, HMGA2 |
| **apoptotic signaling pathway** | 0.007 | 12 | 0.785 | 4 | TRAF1, E2F1, ICAM1, AR, TNF, BOK, IFNB1, CHAC1, IL1B, THBS1, HELLS, IL1A | CSF2, NUPR1, IFNG, BDKRB2 |
| **regulation of apoptotic signaling pathway** | 0.001 | 11 | 0.935 | 2 | TRAF1, E2F1, ICAM1, AR, TNF, BOK, IFNB1, IL1B, THBS1, HELLS, IL1A | CSF2, BDKRB2 |
| **cell cycle process** | 0.000 | 26 | 0.989 | 5 | E2F1, E2F7, EZH2, PTTG1, GPR3, FAM83D, OIP5, NCAPG, IL1B, THBS1, TUBB1, HELLS, IL1A, MKI67, PTTG3P, TPX2, NDC80, BIRC5, GPR132, MCM2, CDC25C, FSD1, WDR62, SPAG5, BUB1B, GADD45A | STOX1, CYP27B1, EPGN, IFNG, HMGA2 |
| **mitotic cell cycle** | 0.000 | 22 | 0.993 | 3 | E2F1, MKI67, E2F7, EZH2, TPX2, BIRC5, NDC80, GPR132, PTTG1, MCM2, CDC25C, FSD1, FAM83D, OIP5, NCAPG, SPAG5, WDR62, BUB1B, IL1B, GADD45A, HELLS, IL1A | STOX1, EPGN, HMGA2 |
| **regulation of cell cycle process** | 0.000 | 16 | 0.927 | 3 | E2F1, MKI67, E2F7, EZH2, TPX2, NDC80, BIRC5, GPR132, CDC25C, GPR3, FAM83D, SPAG5, IL1B, BUB1B, GADD45A, IL1A | STOX1, EPGN, HMGA2 |
| **regulation of mitotic cell cycle** | 0.000 | 14 | 0.866 | 3 | E2F1, MKI67, E2F7, EZH2, TPX2, NDC80, GPR132, BIRC5, CDC25C, IL1B, BUB1B, SIK1, GADD45A, IL1A | STOX1, EPGN, HMGA2 |
| **mitotic nuclear division** | 0.000 | 14 | 0.813 | 3 | TPX2, BIRC5, NDC80, PTTG1, CDC25C, FSD1, FAM83D, NCAPG, SPAG5, OIP5, IL1B, BUB1B, IL1A, HELLS | STOX1, EPGN, HMGA2 |
| **positive regulation of cell proliferation** | 0.141 | 11 | 0.003 | 15 | E2F1, AR, IL6, TNF, NRARP, F3, IL1B, BIRC5, NR4A3, THBS1, FOSL1 | PLAG1, CSF3, CSF2, STOX1, FGFR4, HMGA2, DDR2, ADCYAP1, ALDH3A1, VCAM1, AKR1C2, HOXA3, EPGN, IFNG, MPL |

| | | | | | | |
|---|---|---|---|---|---|---|
| **mitotic cell cycle phase transition** | 0.008 | 11 | 0.975 | 2 | E2F1, E2F7, EZH2, TPX2, BUB1B, GPR132, BIRC5, NDC80, MCM2, CDC25C, GADD45A | STOX1, HMGA2 |
| **positive regulation of cell cycle** | 0.000 | 11 | 0.431 | 4 | FAM83D, E2F1, E2F7, IL1B, BIRC5, NDC80, NR4A3, CDC25C, GADD45A, FOSL1, IL1A | STOX1, EPGN, LRP6, HMGA2 |
| **regulation of cell cycle phase transition** | 0.002 | 10 | 0.910 | 2 | FAM83D, E2F1, E2F7, EZH2, BUB1B, GPR132, BIRC5, NDC80, CDC25C, GADD45A | STOX1, HMGA2 |
| **single-organism organelle organization** | 0.002 | 25 | 0.997 | 5 | E2F1, ICAM1, TNF, TCAP, MYO1B, MAP1A, TPX2, RPH3AL, NDC80, BIRC5, SYNPO2, PACSIN1, PKP1, FRMD7, KRT17, BOK, WDR62, NCAPG, SPAG5, CCDC114, SPTBN2, BUB1B, TUBB1, GADD45A, KATNAL2 | CSF3, NKD2, RND1, CCDC151, MNS1 |
| **cytoskeleton organization** | 0.000 | 22 | 0.827 | 7 | ICAM1, PALM, TNF, TCAP, MYO1B, MAP1A, TPX2, NDC80, BIRC5, SYNPO2, PACSIN1, PKP1, FRMD7, KRT17, ZNF135, SPAG5, WDR62, CCDC114, SPTBN2, TUBB1, GADD45A, KATNAL2 | CSF3, ARC, CCL13, RND1, ZNF135, CCDC151, SIGLEC15 |
| **organelle fission** | 0.000 | 17 | 0.938 | 3 | MKI67, PTTG3P, TPX2, NDC80, BIRC5, PTTG1, CDC25C, GPR3, FSD1, FAM83D, NCAPG, SPAG5, OIP5, IL1B, BUB1B, IL1A, HELLS | STOX1, EPGN, HMGA2 |
| **microtubule cytoskeleton organization** | 0.007 | 10 | 1.000 | 1 | WDR62, SPAG5, CCDC114, MAP1A, TPX2, BIRC5, NDC80, TUBB1, KATNAL2, GADD45A | CCDC151 |
| **response to cytokine** | 0.000 | 20 | 0.114 | 10 | TRAF1, ICAM1, IL6, TNF, IL1RL1, IL1RL2, CXCL3, DUOX2, CXCL2, MCM2, CCRL2, CCL20, TRIM68, IFNB1, F3, TNFRSF19, IL1B, THBS1, FOSL1, IL1A | VCAM1, CSF3, CCL13, GNAO1, CYP27B1, ROBO1, IFNG, TNFSF15, MPL, SELE |
| **inflammatory response** | 0.000 | 16 | 0.082 | 9 | ICAM1, IL6, TNF, IL1RL1, IL1RL2, CXCL3, CXCL2, NLRP3, CCRL2, CCL20, F3, RASGRP1, IL1B, WFDC1, THBS1, IL1A | VCAM1, CCL13, UCN, NUPR1, EPHX2, BDKRB2, PLA2G2D, SELE, ADCYAP1 |
| **positive regulation of cytokine production** | 0.000 | 13 | 0.291 | 5 | IL6, TNF, IL1RL1, IL1RL2, NR4A3, NLRP3, CCL20, RASGRP1, IL1B, THBS1, NLRP10, IL1A, SCAMP5 | CSF2, UCN, EPX, IFNG, ADCYAP1 |
| **single organismal cell-cell adhesion** | 0.005 | 14 | 0.141 | 9 | ICAM1, IL6, TNF, IL1RL2, NR4A3, NLRP3, MYL9, PKP1, IFNB1, NRARP, TNR, RASGRP1, IL1B, ITGA2B | VCAM1, ANXA9, BVES, TENM2, IFNG, LRP6, MPL, PLA2G2D, SELE |
| **chemotaxis** | 0.002 | 13 | 0.551 | 5 | CCRL2, IL6, CCL20, F3, TNR, CXCL3, CXCL2, SPTBN2, IL1B, NR4A3, THBS1, PTGDR2, FOSL1 | VCAM1, CCL13, ROBO1, TENM2, IFNG |
| **leukocyte migration** | 0.001 | 11 | 0.275 | 5 | ICAM1, IL6, TNF, CCL20, CXCL3, CXCL2, IL1B, THBS1, NLRP10, IL1A, ITGA2B | VCAM1, CCL13, EPX, IFNG, SELE |

| | | | | | | |
|---|---|---|---|---|---|---|
| **positive regulation of cellular component movement** | 0.002 | 11 | 0.801 | 3 | ICAM1, IL6, TNF, CCL20, F3, CXCL3, CXCL2, THBS1, MYLK, IL1A, ITGA2B | WNT5B, IFNG, DDR2 |
| **positive regulation of cell migration** | 0.001 | 11 | 0.772 | 3 | ICAM1, IL6, TNF, CCL20, F3, CXCL3, CXCL2, THBS1, MYLK, IL1A, ITGA2B | WNT5B, IFNG, DDR2 |
| **positive regulation of cell motility** | 0.002 | 11 | 0.789 | 3 | ICAM1, IL6, TNF, CCL20, F3, CXCL3, CXCL2, THBS1, MYLK, IL1A, ITGA2B | WNT5B, IFNG, DDR2 |
| **regulation of cell-cell adhesion** | 0.004 | 10 | 0.761 | 3 | IL6, TNF, NRARP, IFNB1, TNR, IL1RL2, RASGRP1, IL1B, NR4A3, NLRP3 | VCAM1, IFNG, PLA2G2D |
| **ion transport** | 0.500 | 13 | 0.003 | 21 | C15ORF48, CPT1B, ICAM1, SPINK1, KCNJ3, PKD2L2, JSRP1, GRIN2C, IL1B, THBS1, SLCO5A1, SIK1, MYLK | C15ORF48, KCNMA1, ARC, SLC38A3, UCN, HNF1A, GNAO1, CACHD1, SLC6A12, BDKRB2, ASIC1, PKD2L1, ADCYAP1, SLC1A2, KCNQ3, CYP27B1, SLC7A3, JSRP1, MCOLN2, PLA2G2D, AKR1C1 |
| **nitrogen compound transport** | 0.692 | 6 | 0.001 | 15 | ABCB9, IL6, TNF, IL1B, BIRC5, UPK3A | CSF2, SLC38A3, UCN, HNF1A, SLC6A12, NUP62CL, IGF2BP1, UPK3A, HMGA2, AMN, AZIN2, ADCYAP1, SLC1A2, SLC7A3, IFNG |
| **response to lipid** | 0.000 | 18 | 0.001 | 16 | E2F1, ICAM1, AR, IL6, TNF, CXCL3, CXCL2, EZH2, NR4A3, NLRP3, NOCT, CCL20, TRIM68, IFNB1, IL1B, WFDC1, THBS1, FOSL1 | CSF3, CSF2, FGFR4, UCN, WNT5B, RBP1, HMGA2, ALDH3A1, ADCYAP1, VCAM1, SSTR2, CYP27B1, HEY1, IFNG, LRP6, SELE |
